# A Generalised Epigenetic Clock Reveals Therapeutic Vulnerabilities Linked to Ageing in Cancer Cells

**DOI:** 10.64898/2026.06.22.733689

**Authors:** Irene Fernández-Rebollo, Alessandro Digilio, Athanasios Oikonomou, Lucia Trastulla, Manel Esteller, Francesco Iorio

**Affiliations:** Computational Biology Research Centre, Human Technopole, Milan, Italy; Università degli Studi di Milano (UNIMI), Milan, Italy; Cancer Epigenetics Group, Sant Pau Research Institute (IRSP), Badalona, Barcelona, Catalonia, Spain; Centro de Investigacion Biomedica en Red Cancer (CIBERONC), Madrid, Spain; Institucio Catalana de Recerca i Estudis Avançats (ICREA), Barcelona, Catalonia, Spain; Physiological Sciences Department, School of Medicine and Health Sciences, University of Barcelona (UB), Barcelona, Catalonia, Spain

**Author notes:** Equally contributing authors (Irene Fernández-Rebollo, Alessandro Digilio).

## Abstract

Epigenetic clocks estimate biological age from DNA methylation patterns but perform poorly in cancer due to extensive epigenetic reprogramming, limiting the study of ageing in tumour biology. Here, we develop GepiClock, an epigenetic clock trained on DNA methylation data from 32 cancer types in The Cancer Genome Atlas. Based on 4,862 CpG sites, GepiClock accurately predicts age across both tumour and normal samples, indicating that core ageing-associated methylation programmes remain detectable despite malignant transformation. Applying GepiClock to molecularly profiled cancer cell lines with matched drug response and CRISPR screening data revealed age-associated vulnerabilities. Younger-predicted cell lines were more sensitive to mTOR, MEK1/2 and HSP90 inhibitors, whereas older lines showed increased sensitivity to AKT and PI3Kα inhibitors. Additional cancer-type-specific patterns and age-associated genetic dependencies were identified. These findings establish a framework to quantify biological age in cancer and link ageing-associated states to therapeutic vulnerabilities.

## Introduction

Over the past century, the average human lifespan has significantly increased, drawing greater focus on ageing and the factors contributing to age-related health decline and diseases^1^. In contrast to chronological age, which merely records the number of years lived, biological age provides a more accurate measure of an individual’s physiological condition and overall health status^2^. It captures the cumulative influence of genetic, environmental, and lifestyle factors on the body’s systems, offering a more integrative reflection of the ageing process.

The search for reliable markers of biological age, distinct from chronological age, has been ongoing for decades. Advances in molecular biology have yielded numerous candidate biomarkers, including telomere length, composite physiological measures, and omics-based estimators derived from transcriptomic, proteomic, and metabolomic data. Among these, epigenetic modifications, particularly DNA methylation patterns, have emerged as the most robust and predictive indicators of biological ageing^3^.

Studies have shown that only a small fraction of Cytosine–phosphate–Guanine (CpG) sites, approximately 2% out of the millions of cytosines in the genome, undergo age-associated changes across tissues in both humans and mice, highlighting the specificity of these epigenetic alterations^4^. Among these, DNA methylation levels at selected CpG sites display a strong and reproducible correlation with age, forming the basis for the development of “epigenetic clocks” (epiclocks): mathematical models that can accurately predict chronological age from DNA methylation patterns^5^. These models have rapidly become indispensable tools in ageing research, offering insights into molecular processes of ageing that chronological age alone cannot capture^6^. The most widely used epiclocks, developed by Horvath in 2013, was trained on DNA methylation data from a wide range of healthy human tissues and cell types. It relies on 353 CpG sites and predicts chronological age with a mean absolute deviation of 3.6 years^7^. Around the same time, Hannum and colleagues developed a blood-based clock built on 71 CpG sites, which showed a similarly strong correlation with age and a mean absolute deviation of 4.9 years^8^. In 2018, Horvath introduced an updated clock incorporating 391 CpGs, designed to improve prediction accuracy across a broader spectrum of tissues and cell types^9^. More recently, second-generation clocks have been developed, such as Levine’s PhenoAge, which integrates measures of morbidity and mortality into the model and improves predictive power for health outcomes beyond that of the first generation^10^. Although current epiclocks accurately predict age in healthy samples, they perform poorly in cancer contexts due to the distinct epigenetic alterations that characterise malignancy. As a result, they inconsistently estimate patients’ chronological age and fail to reliably capture their biological ageing state^7,8^. This limitation reflects the fact that age-related methylation signatures are obscured by the widespread and aberrant DNA methylation changes that play a central role in cancer development and progression^11,12^.

Indeed, most human tumours exhibit imbalances in DNA methylation patterns, with three significant alterations commonly observed: a global loss of DNA methylation^13^, increased levels of DNMT1^14^, and regional imbalances in DNA methylation at CpG islands, leading to the silencing of tumour suppressor genes and the activation of oncogenes^15^. In addition, cancer cells more often exhibit epigenetic drift characterised by stochastic processes that involve both gains and losses of the CpGs methylation state^16^, resulting in their methylation patterns changing unpredictably compared to healthy cells. Consequently, understanding and identifying the functional consequences of these patterns is essential for improving tumour diagnosis, prognosis, and treatment^17^. More broadly, cancer and ageing can be seen as two different manifestations of the same basic process, the accumulation of cellular damage over time, often sharing common mechanistic foundations known as the “hallmarks”^18^, among which epigenomic alterations play a central role^19^. However, studying ageing in the context of cancer remains particularly challenging, as widespread epigenetic reprogramming obscures canonical age-associated methylation signatures.

Despite these challenges, the wealth of publicly available multi-omic resources offers an unprecedented opportunity to investigate how ageing-related mechanisms intersect with cancer vulnerabilities. Thousands of immortalised cancer cell lines (CCLs) have been deeply profiled at the genomic, transcriptomic, epigenomic, and proteomic levels, and systematically tested for their responses to large panels of drugs and genetic perturbations^20–24^. Yet this rich landscape remains underexploited from the perspective of ageing biology. A key limitation is that the chronological age of a cancer cell line is inherently ill-defined and biologically uninformative, as it cannot be meaningfully inferred from donor age, time in culture, or passage number, nor does it reflect the cumulative epigenetic drift and adaptive reprogramming that occur during tumour evolution and long-term in vitro propagation^25–27^. A further complication is that CCLs often diverge substantially from the tumours they are intended to model. This affects multiple molecular layers, including the genome, transcriptome, and particularly the methylome, due to prolonged culture, selective pressures, and stochastic epigenetic drift^28^. As a result, the DNA methylation profiles of many CCLs bear little resemblance to those of their parental tumours, raising concerns about their fidelity as cancer models for epigenetic studies^29–31^. Nevertheless, the epigenetic age of a CCL may still capture biologically meaningful methylation signatures that reflect aspects of cellular ageing (even if partially) and could serve as biomarkers of drug response or genetic dependency.

Here, we present the generalised epigenetic clock (GepiClock): an epigenetic clock specifically trained on DNA methylation data from tumour samples, which generalises across both cancer and healthy contexts. By modelling cancer settings along a gradient of age-related phenotypes, our generalised clock not only achieves high predictive performance but also provides a robust measure of biological age that can be applied to CCLs, despite their complex and aberrant methylomes. Most importantly, our model enables the incorporation of biological age as a feature in the analysis of multi-omic and clinical data, allowing systematic association of ageing signatures with drug responses and genetic dependencies. Through this framework, we reveal pan-cancer as well as tumour-specific associations between predicted age and drug-response/genetic-dependencies, highlighting previously unreported mechanisms of age-related tumour biology and uncovering therapeutic opportunities that are dependent on the ageing state of cancer cells.

## Results

### A generalised epigenetic clock efficiently predicts biological age in cancer and healthy samples

We considered DNA methylation data of tumor samples from the Cancer Genome Atlas (TCGA)^32^, encompassing 32 different cancer types and a wide range of chronological ages (**Supplementary Fig. 1-2** and **Supplementary Table 1**).

To enable robust application across methylation platforms, we focused on the 369,845 CpGs shared between Illumina 450K, EPIC, and EPIC v2.0 arrays (**Supplementary Fig. 3**).

Consistently with previously proposed clocks, such as the Horvath 2013 Clock^7^, we developed a generalised epigenetic clock (GepiClock) as an elastic net regression model (**Methods**), trained exclusively on primary tumour samples after excluding all samples derived from patients with matched normal tissue data (**Fig. 1A**). We analysed a total of 9,105 TCGA samples splitting them into training (80%) and test (20%) sets using stratified sampling by cancer type and age, with 10-fold cross-validation applied. The resulting model comprised 4,862 variables with associated non null regressor coefficients, i.e. the predictive methylation probes, or CpGs (**Supplementary Table 2**). We evaluated model performance on the held-out test set, as well as on an independent set of 733 TCGA patient samples not included in the train-test split, for which both matched tumour and normal tissues were available (**Fig. 1A**). In the independent evaluation set, the GepiClock predicted age showed a Pearson correlation exceeding 0.76 with chronological age when considering tumour samples and an even higher correlation of 0.85 for matched normal samples (**Fig. 1B**). When benchmarking predicted-versus-chronological age correlation obtained by the GepiClock against those obtained with established epiclocks on the same evaluation sets, we observed consistently higher scores for our model across both tumour and normal samples (**Fig. 1C**). This performance advantage was consistently maintained when evaluated using mean absolute error (MAE), with GepiClock achieving values of ±6.8 years in tumour samples from the training set, ±7.1 years in tumour samples from the independent test set, and ±7.0 years in normal samples, corresponding to an order-of-magnitude improvement in tumour contexts compared to existing clocks, further supporting the robustness of the model across metrics (**Supplementary Fig. 4** and **supplementary Table 2**).

**Fig. 1.**
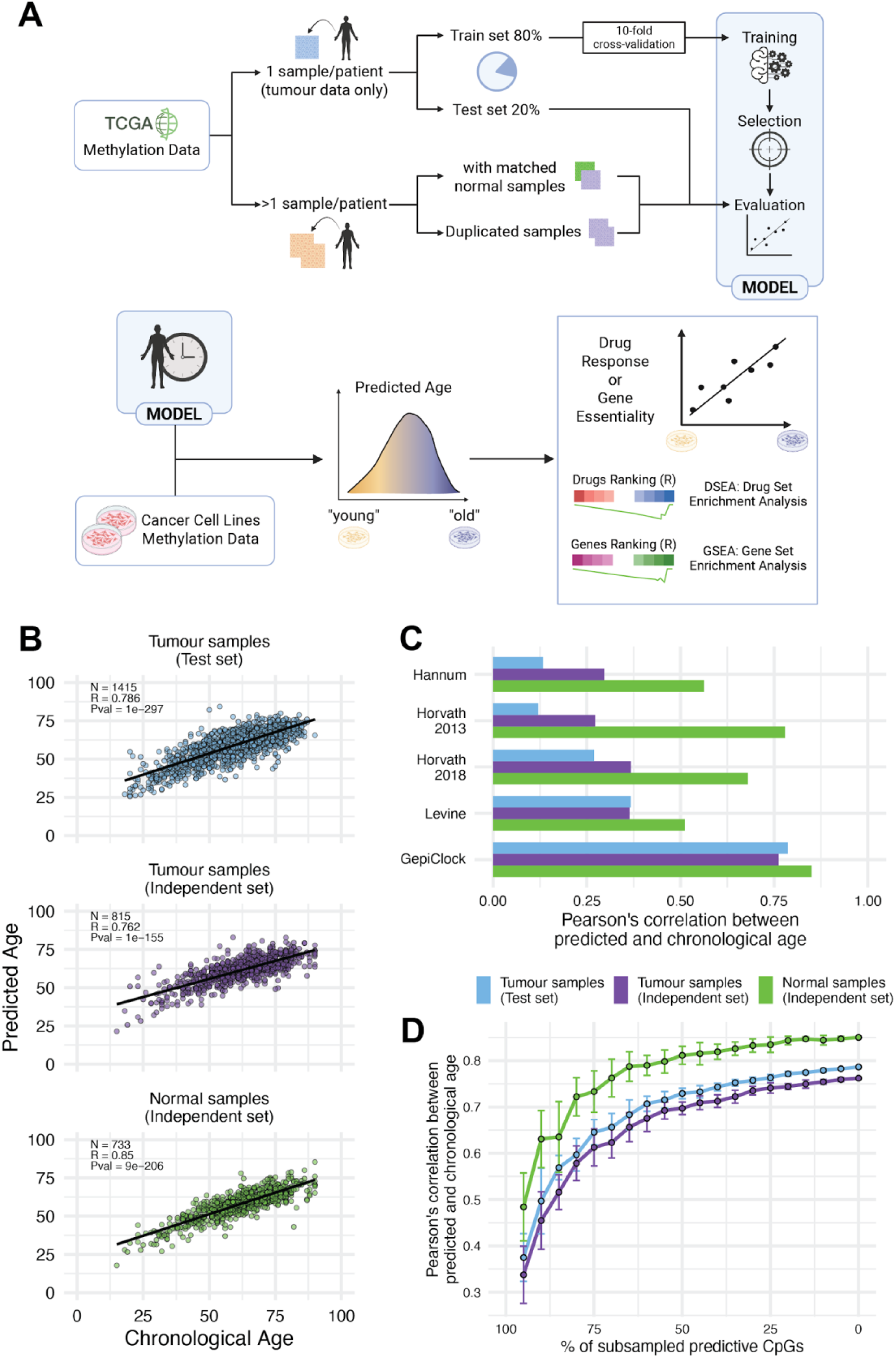
Schematic of the generalised epigenetic clock and performance summary. A. Overview of the generalised epigenetic clock training and application. TCGA primary tumour samples are used for model training, with matched tumour and normal samples reserved for performance evaluation. The model is subsequently applied to cancer cell line methylation data to stratify samples by predicted biological age for downstream analyses of differential drug response and genetic dependency, enabling the identification of vulnerabilities putatively associated with ageing-related cellular processes. Created with BioRender.com. B. Pearson’s correlation (R) of predicted vs chronological age in different TCGA data type subgroups (Tumour samples in test set, tumour samples in the held-out independent set, and normal samples in the held-out independent set). C. Predictive performances (R values) of widely used epiclocks and of the GepiClock (Rebollo) across different data type subgroups (defined as in B). D. GepiClock predictive performances across sample subgroups (as defined in B and C) when keeping in the model fixed percentages of randomly sub-sampled predictive CpGs, across 20 sampling trials. Average values are reported with whiskers indicating 25%-75% percentiles.

We further validated the performance of the GepiClock on an independent external dataset (**Methods**) comprising DNA methylation profiles from 289 women enrolled in the NCI-Maryland Breast Cancer Cohort^33^. This dataset includes 185 tumour samples, 113 matched adjacent normal tissues, and 104 normal breast samples from reduction mammoplasty. Importantly, the cohort spans individuals of African American, European American, and Kenyan ancestry, thereby encompassing populations that are typically underrepresented in large-scale genomic resources such as TCGA.

In this external setting, the GepiClock maintained high Pearson correlations between predicted and chronological age across tumour, normal, and normal-adjacent samples, and overall outperformed previously established clocks. A similar trend was observed in the independent external dataset, where GepiClock achieved MAE values of ±8.3 years in tumour samples, ±8.6 years in adjacent normal tissues, and ±6.4 years in normal samples, consistently outperforming all other clocks, particularly in tumour samples where errors remained substantially lower than those of existing models (**Supplementary Fig. 5** and **supplementary Table 2**).

Given the relatively large number of predictive CpGs in the GepiClock compared with other clocks (**Supplementary Fig. 6**), we assessed its robustness to missing CpG probes and explicitly tested whether performance gains could be attributed solely to increased model complexity. To this end, we evaluated model performance after matching the number of CpG probes to those included as predictive in other clocks by retaining an equivalent number of randomly selected predictive CpGs (**Methods**). This analysis also allowed us to assess the compatibility of the clock in the presence of missing measurements, as commonly encountered when applying the model across different DNA methylation platforms with partially overlapping probe sets. This represents a conservative evaluation, as it penalises the GepiClock by removing predictive CpGs post hoc rather than re-training under fixed model-size constraints. Despite of this, our model maintained, on average, good predictive accuracy even when the predictive CpGs were reduced down to 50% (average Pearson’s correlation = 0.7 and 0.81, respectively for tumour and normal samples, in the independent evaluation set; **Fig. 1D**). Moreover, when restricting the GepiClock to a number of predictive CpGs equal to those used by existing clocks, it continued to substantially outperform them in tumour samples (**Supplementary Fig. 7**). This indicates that our model’s performance is not merely driven by its complexity, in terms of number of regressors. Importantly, this robustness enables reliable application of our model to DNA methylation data generated using different array platforms, where subsets of the predictive CpGs may be absent.

### Characterisation of the GepiClock predictive CpGs and comparison with previous epiClocks

Overall, the GepiClock includes a larger number of predictive CpGs (the GepiClock CpGs) than previously described clocks, reflecting the use of a substantially broader CpG input space compared with other models, most of which are trained on the legacy 27K Illumina Infinium Methylation BeadChip. In addition, training on tumour samples spanning multiple cancer types increased data heterogeneity due to both malignant epigenetic states and tissue diversity. Consequently, the sets of CpGs selected by different clocks exhibit limited overlap (**Supplementary Fig. 6**). However, this mirrors the sparse concordance, in terms of shared predictive CpGs, that is generally observed across all pairwise comparisons of existing epiclocks (**Supplementary Fig. 6**).

At the CpG level, GepiClock shows the greatest similarity to the Horvath 2018 clock, sharing 36 predictive CpGs (respectively 0.74% and 9.21%), the highest overlap observed among all pairwise clock comparisons (**Supplementary Fig. 6**). At the gene level, however, the strongest similarity emerges with the Levine clock (105, genes in common, respectively 3.15% and 17.83%, **Supplementary Fig. 6**). Across all clocks, four CpGs are shared (cg06493994, cg22736354, cg09809672, and cg19722847), mapping to the SCGN, NHLRC1, EDARADD, and IPO8 genes, respectively. In addition, two genes, KLF14 and SFMBT1, are consistently shared across clocks at the gene level. Notably, KLF14 is a transcription factor involved in metabolic regulation, which represents one of the most reproducible loci associated with age-related DNA methylation changes^34^, while SFMBT1 encodes a chromatin regulator linked to Polycomb-mediated repression, a class of genomic regions known to accumulate age-associated methylation^35^. Together, the conservation of these loci across independently derived clocks suggests that a core subset of ageing-associated epigenetic signals remains detectable despite substantial differences in training data and CpG selection strategies. Moreover, GepiClock identifies a largely novel set of CpGs (4,761) and associated genes (3,033) not represented in previous clocks, suggesting that these features capture conserved ageing-associated methylation programmes that persist across both normal and tumour contexts, consistent with the model’s robust performance in both settings (**Supplementary Fig. 6**).

To characterise the genomic context of the GepiClock CpGs, we examined their distribution across autosomal chromosomes. We observed a nearly uniform distribution of GepiClock CpGs across mapping chromosomes, over the total number of Illumina Array mapped on each chromosome (normalised Shannon’s entropy = 0.96, average ratio across chromosomes = 4.54, with standard deviation = 2.18, **Fig. 2A**). The highest proportion of CpGs selected by the model as predictive (the GepiClock CpGs) fall in chromosomes 1, 6 and 19 (with many regions significantly enriched of predictive CpGs), while chromosomes 18, 20, 21 and22 are the least represented (**Fig. 2A, left panel**); this roughly reflects the chromosome representation among the CpGs in Illumina arrays, thus the lack of significant biological or technical biases (**Supplementary Fig. 8**).

**Fig. 2.**
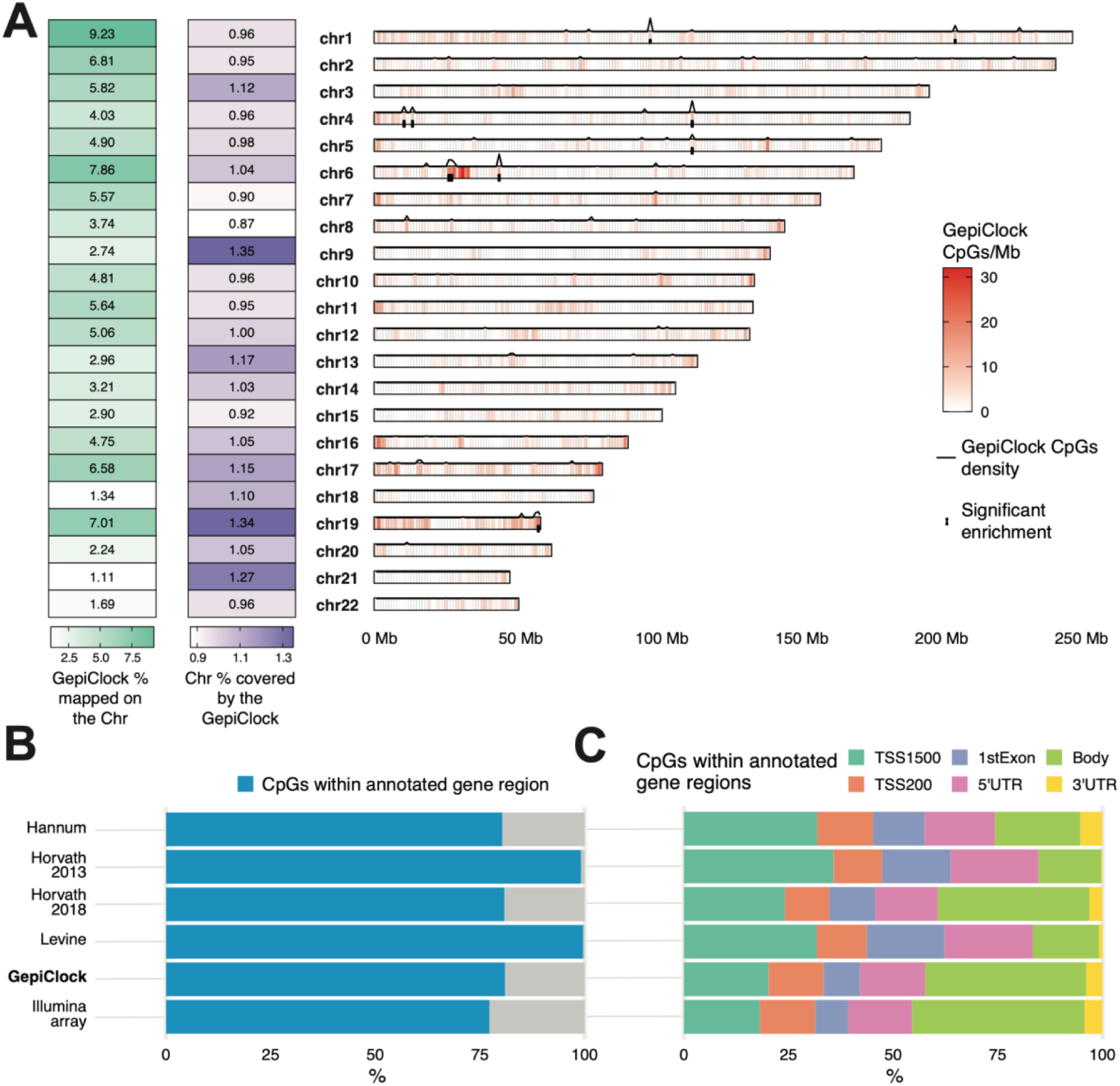
Genomic distribution of the GepiClock predictive CpGs. A. Distribution of the GepiClock CpGs across autosomal chromosomes. On the left: percentages of GepiClock CpGs mapped onto each chromosome; In the centre: percentages of all possible CpGs mapped to each chromosome that are included in the GepiClock as predictive. On the right: density distributions of predictive CpGs (CpGs/Mb) along chromosomes. For each 1Mb genomic window, we compared the observed number of predictive CpGs to the expected number based on the total CpG representation in the array through a hypergeometric test. The black line above each chromosome represents the log2 of the resulting p-value corrected for multiple hypothesis testing, and black squares indicate windows in which predictive CpGs are significantly enriched (corrected hypergeometric p-value <= 0.05) relative to random expectation. BC. Proportion of CpGs with annotated gene region (B) and their functional genomic annotation within gene regions (C) across epiclocks and the Illumina array (only probes shared among the different Infinium BeadChips are considered).

We next compared the different epiclocks in terms of functional genomic annotation. The proportion of GepiClock CpGs mapping to annotated gene regions was comparable to that observed for the Hannum and Horvath 2018 clocks, as well as to the distribution of CpG probes shared across Illumina Infinium methylation arrays (Shannon-Jensen (SJD) distance = 4.08e-05, 2.54e-06 and 0.001, respectively, **Fig. 2B**, and **Supplementary Fig. 9A**).

We further examined the localisation of these CpGs within gene features and observed a broadly similar distribution across all epiclocks, including promoter regions (TSS1500, TSS200, 5′UTR, and first exon), gene bodies, and 3′UTRs (average SJD = 0.026, **Supplementary Fig. 9A**). Overall, the GepiClock closely follows the genomic distribution of the Illumina array, with a modest reduction, compared to other epiclocks, in the proportion of CpGs located in promoter regions (**Fig. 2C** and **Supplementary Fig. 9C**).

To functionally characterise the biological context of the GepiClock CpGs, we performed a functional enrichment analysis of the genes in their nearly located sites. Among the significantly enriched functional classes (adjusted hypergeometric test p-value < 0.05) we observed terms involved in developmental and morphogenetic processes, including embryonic organ development, pattern specification, epithelial tube morphogenesis, and skeletal system development (**Supplementary Fig. 10** and **Supplementary Table 3**). These enrichments were characterised by a significant representation of WNT and BMP pathway components, key regulators of cell fate and tissue patterning^36,37^, pointing to developmental programmes that remain active in adulthood and are progressively remodelled during ageing. Additional enriched terms highlighted processes related to cell adhesion and cytoskeletal organisation, such as homophilic cell adhesion and actin filament organisation (**Supplementary Fig. 10** and **Supplementary Table 3**). Together, these results indicate that CpG sites predictive of chronological age preferentially localise near developmental regulators and pathways governing cell identity and tissue architecture, biological systems that are frequently implicated in both ageing and tumorigenesis.

### Predicted Age as a potential biomarker of drug response and genetic dependencies in cancer cells

We next asked whether the biological age predicted by the GepiClock could serve as a meaningful molecular descriptor of cancer cell lines (CCLs) and stratify their therapeutic vulnerabilities. Importantly, our objective was not to predict the chronological age of CCLs, which is intrinsically ill-defined and biologically ambiguous, as it cannot be meaningfully derived from donor age, time since derivation, or passage number. Consistently, the GepiClock does not reliably recapitulate the annotated age of CCLs (**Supplementary Fig. 11**), nor would such a comparison be biologically informative. Moreover, it is well established that CCLs undergo extensive epigenetic drift and culture-induced reprogramming, resulting in methylation landscapes that substantially diverge from those of their parental tumours^27,29,31,38,27,29,38^. Rather than treating this divergence as a limitation, we leveraged the GepiClock to project CCL methylation profiles onto an ageing-relevant epigenetic axis derived from primary tumours, thereby capturing conserved ageing-associated methylation signatures that persist despite broader epigenomic disruption. This approach enables stratification of CCLs according to their epigenetic similarity to tumour-associated ageing states. Importantly, this strategy is conceptually distinct from pharmacoepigenomic approaches that identify locus-specific methylation biomarkers predictive of individual drug responses^39^. Instead of modelling direct associations between CpG methylation levels and drug sensitivity, we infer a higher-order biological state reflecting ageing-related epigenomic organisation and interrogate how this latent state relates to pharmacological and genetic vulnerabilities. As such, our framework complements existing epigenetic biomarker strategies by providing a global, ageing-informed lens through which cancer dependencies can be interpreted.

To investigate whether this ageing-informed stratification might associate with differential drug response and genetic vulnerabilities, we performed comprehensive analyses in both pan-cancer and cancer type-specific settings (**Methods**, **Supplementary Fig. 12** and **13**) using publicly available DNA methylation data from a large panel of cancer cell line models that we previously characterised^23^.

We integrated these data with publicly available drug response measurements from systematic screens conducted on the same cell lines by the Genomics of Drug Sensitivity in Cancer (GDSC) database^40^. To isolate the contribution of predicted biological age to drug sensitivity, we fitted an analysis of variance (ANOVA) model (**Methods**) that accounts for variability attributable to tissue of origin, cancer type, and microsatellite instability (MSI) status. We then computed Pearson correlation coefficients (R) between predicted biological age and the residual ln(IC50) values for each drug, applying false discovery rate (FDR) correction to control for multiple hypothesis testing (**Methods**).

In the pan-cancer setting, we found 69 significant (adjusted p-value < 0.05) yet overall modest (−0.17 < R < 0.32) correlations (**Supplementary Fig. 14** and **Supplementary Table 4**).

Given the modest effect sizes observed at the level of individual compounds, we next asked whether coherent patterns might nevertheless emerge at the level of drug target classes. We therefore performed a Drug Set Enrichment Analysis (DSEA), ranking compounds according to their Pearson correlation coefficient between predicted biological age and residual ln(IC50) values (**Methods**). This approach enabled us to test whether drugs targeting specific molecular pathways were systematically enriched among those showing positive or negative age-associated correlations.

DSEA revealed significant (DSEA adjusted p < 0.05) positive enrichment for mTOR, MEK1/2, and HSP90 inhibitors, and significant negative enrichment for AKT1/2/3 and PI3Kα inhibitors (**Fig. 3A, Supplementary Table 5**). Per construction, positive enrichment indicates that higher predicted biological age is associated with increased ln(IC50) values, that is, reduced drug sensitivity, whereas negative enrichment indicates that higher predicted age corresponds to lower ln(IC50) values and therefore greater sensitivity. Accordingly, cell lines predicted as biologically younger display increased sensitivity to mTOR, MEK1/2, and HSP90 inhibitors, while biologically older cell lines exhibit enhanced sensitivity to AKT and PI3Kα inhibitors (**Fig. 3A, Supplementary Table 5**). These findings indicate that, although individual drug-level effects are modest, predicted biological age captures coordinated sensitivity patterns across functionally related therapeutic classes.

**Fig. 3.**
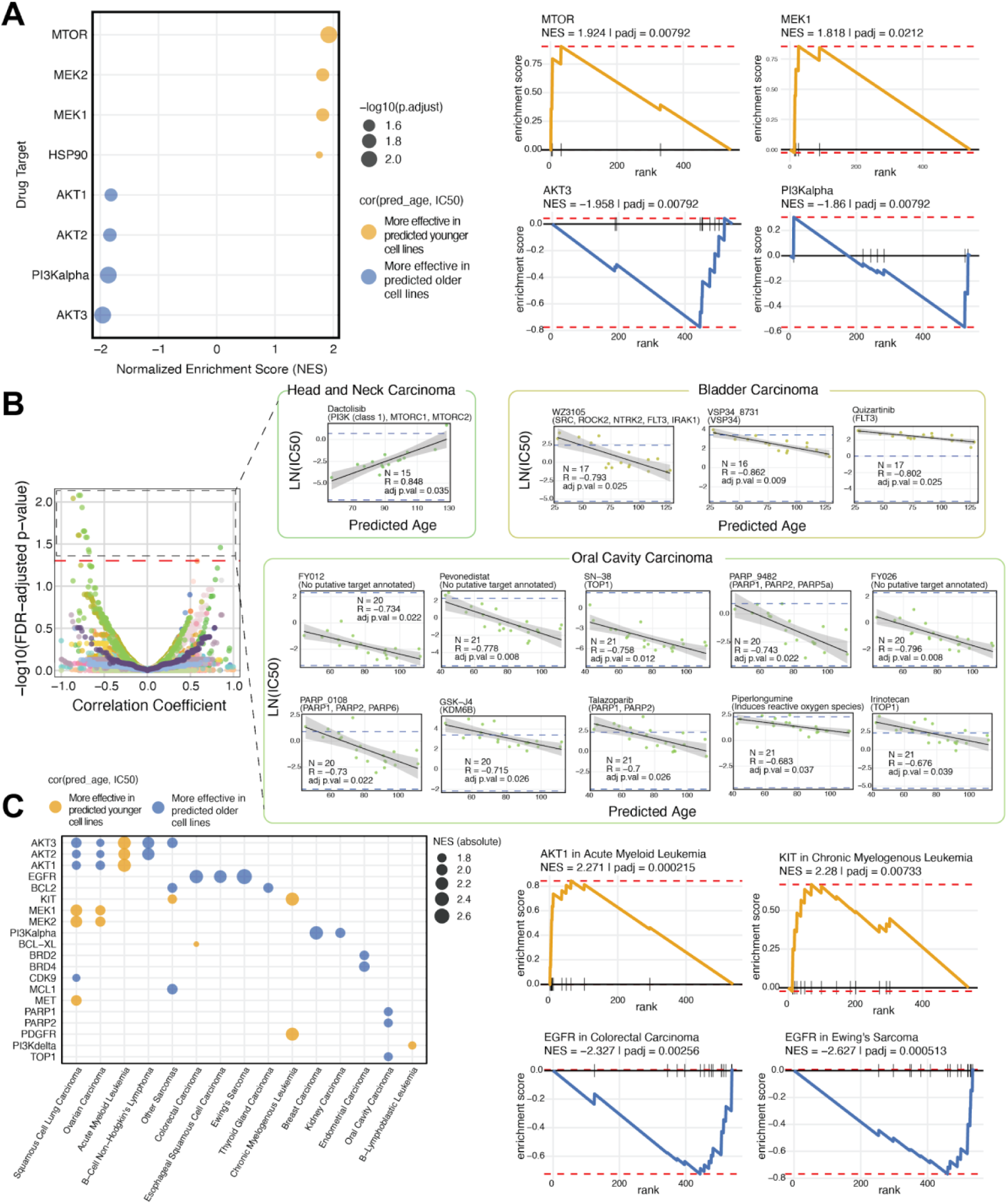
Predicted biological age as a molecular signature of drug response in pan-cancer and cancer type–specific analyses. A. Pan-cancer Drug Set Enrichment Analysis (DSEA). Drugs were ranked according to the Pearson correlation between predicted age and drug sensitivity (ln(IC50)). Left: enriched drug target classes are shown on the y-axis with their corresponding normalized enrichment scores (NES) on the x-axis. Dot size represents statistical significance (-log10 adjusted p-value), while colour indicates the direction of enrichment: positive NES (yellow) corresponds to drugs for which increased predicted age is associated with higher IC50 (reduced sensitivity), whereas negative NES (blue) corresponds to drugs for which increased predicted age is associated with lower IC50 (increased sensitivity). Right: representative enrichment plots for the two most positively enriched target classes (MTOR, MEK1) and the two most negatively ones (AKT3, PI3Kα). In each plot, the curve represents the running enrichment score across the ranked drug list, and vertical ticks indicate drugs belonging to the corresponding target class. B. Individual drug associations. Left: volcano plot showing correlation coefficients between predicted age and ln(IC50) for all drug-cancer type pairs (x-axis) and corresponding -log10(FDR-adjusted p-values) (y-axis). Right: scatterplots for drug-cancer type pairs with statistically significant associations. Each point represents a cell line plotted by predicted age (x-axis) and ln(IC50) (y-axis). Shaded areas indicate the confidence interval of the regression line, and dashed blue lines mark the experimentally tested drug concentration ranges. Plot titles report the compound name together with its nominal target annotation. C. Cancer type-specific DSEA. Drug set enrichment analyses were performed independently for each cancer type by ranking compounds according to the correlation between predicted age and ln(IC50). Each dot represents a significantly enriched drug target within a given cancer type. Dot colour indicates the direction of enrichment (yellow: higher IC50 with increasing predicted age; blue: lower IC50 with increasing predicted age), while dot size reflects the absolute NES. Enrichment plots for representative significantly positive and negative targets enriched are shown.

Notably, these target classes broadly map onto nutrient-sensing, growth, and proteostasis pathways that have been extensively implicated in ageing biology^41–43^, suggesting that the ageing-associated epigenetic state captured by GepiClock may reflect differential reliance on proliferative versus stress-adaptive signalling programmes.

As ageing-related epigenetic states may interact with lineage-specific oncogenic programs, we next evaluated associations within individual cancer types.

In the cancer type-specific setting, 780 cell-lines could be included in the analysis (**Methods**, **Supplementary Fig. 12-13**). Within each cancer type, we quantified the association between predicted biological age and drug response by computing Pearson correlations with ln(IC50) values. When microsatellite instability (MSI) subgroups were present, we adjusted ln(IC50) values using an ANOVA model and correlated predicted age with the resulting residuals; in cancer types without MSI variability, correlations were computed directly on ln(IC50) values (**Supplementary Fig. 14**).

We observed statistically significant associations (adjusted p < 0.05) between predicted biological age and individual drug responses in three cancer types (**Fig. 3B** and **Supplementary Table 4**). In Head and Neck Squamous Cell Carcinoma (HNSC) cell lines, Dactolisib (a PI3K/mTOR dual inhibitor) displayed a strong positive correlation (R = 0.85), indicating reduced sensitivity with increasing predicted age. In contrast, Bladder Carcinoma CCLs exhibited strong negative correlations (increased sensitivity for predicted older CCLs) for VSP34-8731 (a PIK3C3 inhibitor), Quizartinib (a FLT3 inhibitor), and WZ3105 (an EGFR inhibitor). Similarly, Oral Cavity Carcinoma CCLs showed negative correlations across a broader panel of compounds, including FY026 (CDK inhibitor), Pevonedistat (NAE/NEDD8-activating enzyme inhibitor), SN-38 (topoisomerase I inhibitor), PARP-9482 (PARP1/2 inhibitor), FY012 (CDK inhibitor), PARP-0108 (PARP1/2 inhibitor), GSK-J4 (KDM6A/UTX–KDM6B/JMJD3 histone demethylase inhibitor), Talazoparib (PARP1/2 inhibitor), Piperlongumine (ROS inducer/oxidative stress modulator), and Irinotecan (topoisomerase I inhibitor)(**Fig. 3B**). With the exception of Quizartinib in Bladder Carcinoma and SN-38 in Oral Cavity Carcinoma - where ln(IC50) values were entirely above or within the experimentally tested concentration range, respectively - all significant associations involved drug response estimates spanning both within-range and extrapolated values beyond the highest tested concentrations. This pattern indicates that the observed correlations are unlikely to be artefacts of dose-range saturation and instead reflect biologically meaningful stratification of drug response.

Similarly to the pan-cancer analysis, we performed a DSEA independently within each cancer type by ranking drugs according to their age-associated correlation coefficients. Several tumour types exhibited significant enrichment (DSEA adjusted p < 0.05) of specific drug target classes (**Fig. 3C**, **Supplementary Table 5**). Recurrent patterns emerged across multiple cancer types, including EGFR, PI3Kα, and BCL2 inhibitors associated with increased sensitivity in predicted older cell lines, and KIT and MEK1/2 inhibitors showing greater efficacy in predicted younger cell lines. These recurrent enrichments are consistent with a shift from growth factor-driven dependencies in predicted younger states to survival and apoptotic signalling vulnerabilities in predicted older states.

A notable pattern was observed for AKT1/2/3 inhibitors, which exhibited increased efficacy in predicted older cell lines from Squamous Cell Lung Carcinoma, Ovarian Carcinoma, Burkitt’s Lymphoma, and Osteosarcoma, whereas the opposite association - enhanced sensitivity in predicted younger cell lines- was observed in Acute Myeloid Leukemia. This context-dependent behaviour may reflect lineage-specific wiring of PI3K-AKT signalling, particularly in haematological malignancies such as Acute Myeloid Leukemia, where AKT activation is often more tightly coupled to proliferative and differentiation programmes than to the stress-adaptive signalling states frequently observed in solid tumours^44,45^. In contrast, other enrichments appeared to be tumour-type specific. For example, BRD2/4 inhibitors in Endometrial Carcinoma and PARP1/2 inhibitors in Oral Cavity Carcinoma were enriched among drugs to which predicted older CCLs were more sensitive, whereas MET inhibitors in Squamous Cell Lung Carcinoma and PDGFR inhibitors in Chronic Myelogenous Leukemia were preferentially effective in predicted younger CCLs (**Fig. 3C**). These lineage-restricted associations suggest that the interaction between ageing-associated epigenetic states and therapeutic dependencies may be modulated by tumour-specific signalling architecture. Notably, the compounds identified within each tumour type converge on biologically coherent pathway themes. In Oral Cavity carcinoma, drugs targeting DNA damage response, replication stress, and chromatin regulation (TOP1, PARP, CDK, and KDM6 inhibitors) were preferentially effective in predicted older cell lines, consistent with increased reliance on genome maintenance and epigenetic regulation programmes. In Bladder Carcinoma, age-associated sensitivity involved receptor tyrosine kinase and autophagy-related pathways (EGFR, FLT3, and VPS34), suggesting differential engagement of survival signalling circuits. In contrast, in HNSC, the association with the dual PI3K/mTOR inhibitor Dactolisib indicates that ageing-associated epigenetic states may interact with lineage-specific growth signalling dependencies (**Supplementary Table 5**). This supports a context-dependent link between ageing-related epigenomic organisation and therapeutic vulnerabilities.

Taken together, these findings indicate that the ageing-associated epigenetic state captured by GepiClock stratifies therapeutic vulnerabilities across both shared and lineage-specific contexts.

We next applied a similar framework to assess whether stratifying CCLs by predicted biological age reveals differential genetic dependencies. Using gene effect scores from the Cancer Dependency Map (DepMap)^22,46^, which quantify gene essentiality as the reduction in cellular viability following CRISPR-mediated gene knockout, we analysed both pan-cancer and cancer type-specific contexts (**Supplementary Fig. 15-16**), treating predicted age as a continuous variable and correlating it with gene dependency values (**Methods**). As observed for drug-response associations, pan-cancer correlations between predicted biological age and gene effect scores were generally modest in magnitude (−0.199 < r < 0.173, **Supplementary Table 6**). To determine whether coordinated biological signals were nevertheless present, we performed a Gene Set Enrichment Analysis (GSEA), ranking genes according to their age-associated correlation coefficients to assess enrichment of functional pathways among those showing significant associations. Specifically, we ranked genes by the Pearson correlation between gene dependency scores and predicted biological age across CCLs and performed GSEA to identify enriched pathways.

Using Gene Ontology Molecular Function (GOMF), Gene Ontology Cellular Component (GOCC), and KEGG pathway annotations, we observed statistically significant results for multiple biological programmes in the cancer-type specific setting (adjusted p < 0.05; **Supplementary Figs. 17-19** and **Supplementary Table 7)**.

At the pan-cancer level, GSEA revealed coherent enrichment patterns despite the modest magnitude of individual gene-level associations. In particular, gene sets associated with transcriptional regulatory activity, including DNA-binding transcription activator activity, were enriched among dependencies correlated with higher predicted biological age (**Supplementary Fig. 17**).

In contrast, gene sets associated with membrane and endoplasmic reticulum (ER) compartments, including the ER membrane and ER subcompartments, were enriched among dependencies associated with predicted younger CCLs (**Supplementary Fig. 18**). Similarly, KEGG pathway analysis highlighted enrichment signalling-related pathways among genes associated with younger predicted states (**Supplementary Fig. 19**).

Together, these patterns suggest a shift from signalling- and membrane-associated processes in predicted younger cell lines toward increased reliance on transcriptional regulatory programmes in older epigenetic states. This transition is consistent with ageing-associated rewiring of cellular regulation, whereby dynamic signalling and environmental responsiveness are progressively replaced by more stable, transcriptionally driven regulatory states.

We also observed several significant enrichments at the individual cancer-type level (**Supplementary Fig. 20-23**). The three most significant terms per cancer type, ranked by Normalized Enrichment Score (NES), are summarised in **Fig. 4A**.

**Fig. 4:**
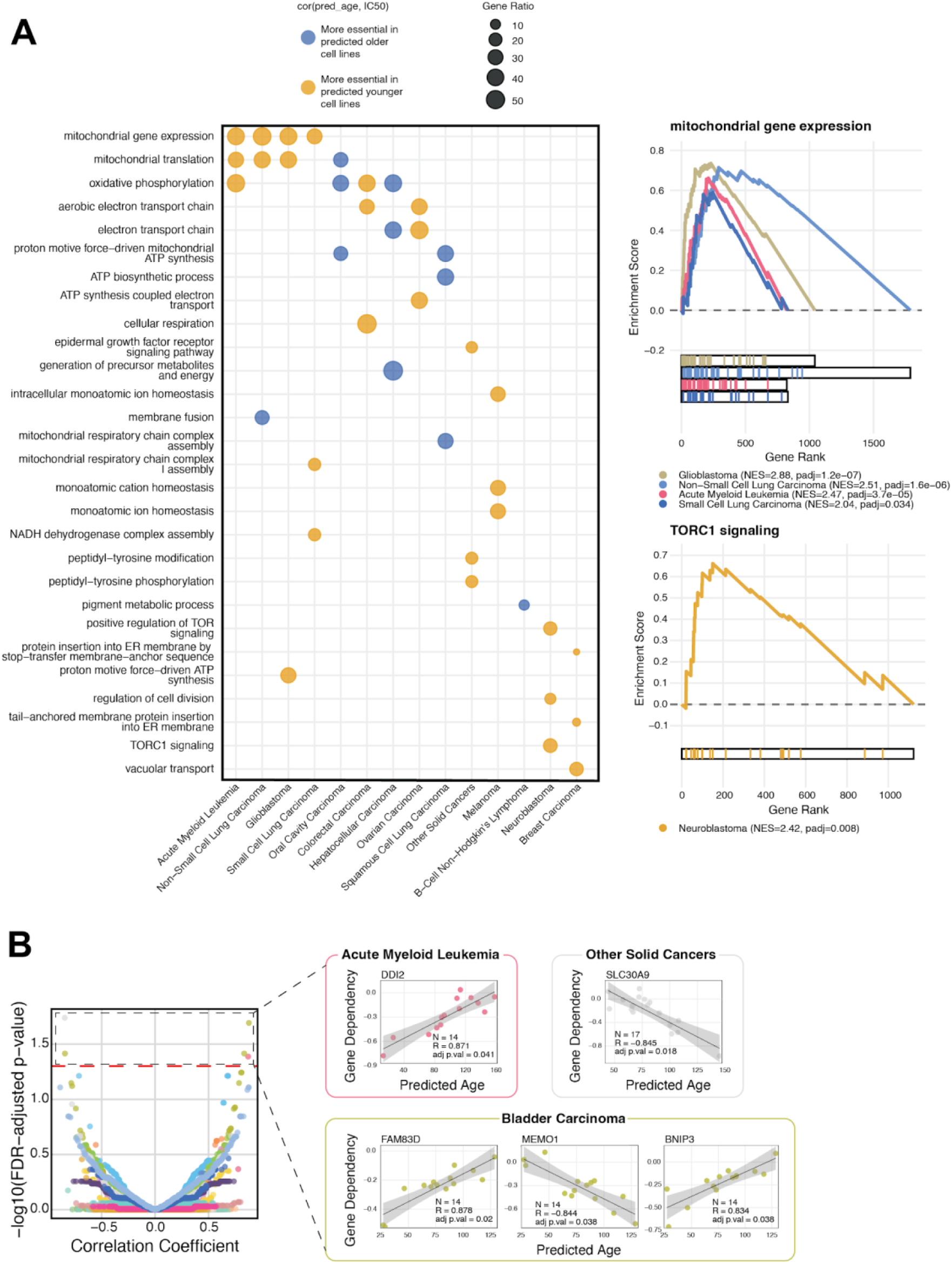
Predicted Age as a potential molecular signature for genetic dependencies in a cancer type-specific setting. A. Cancer type-specific Gene Set Enrichment Analysis (GSEA) performed individually using GOBP terms for each cancer type by ranking genes according to their Pearson’s correlation coefficient between predicted age and gene dependency. For each cancer type, the three most significant terms (y-axis), based on their adjusted p-values, are displayed. Each dot represents a significantly enriched biological process within a specific cancer type (x-axis), with the dot color indicating the direction of the correlation (with yellow as positive, and blue as negative) and the corresponding Normalized Enrichment Score (NES) value, with the dot size reflecting the gene ratio. B. Selected examples of significantly enriched categories. C. The volcano plot (left) shows the correlation coefficients (x-axis) between predicted age and gene dependency and corresponding –log₁₀(FDR-adjusted p-valyes) (y-axis) for the gene–cancer type pair, further highlighting those that show statistically significant correlations with correlation scatterplots (right).

Notably, gene sets related to mitochondrial gene expression and mitochondrial translation were enriched among dependencies associated with predicted younger CCLs across multiple cancer types, suggesting increased reliance on mitochondrial maintenance and proteostasis programmes. In contrast, predicted older CCLs showed enrichment for oxidative phosphorylation, electron transport chain activity, and respiratory processes, particularly in oral cavity, hepatocellular and squamous cell lung carcinoma CCLs. This pattern is consistent with age-associated mitochondrial remodelling, whereby declining respiratory efficiency and accumulated mitochondrial stress are accompanied by increased dependence on mitochondrial maintenance and protein quality-control pathways^47^. The persistence of these signatures in cancer cell lines suggests that ageing-related epigenetic states captured by GepiClock stratify distinct mitochondrial dependency programmes within tumour models, consistent with the continued importance of mitochondrial function for cancer cell survival and metabolic adaptation^48^.

Another notable example included enrichment of membrane fusion among dependencies associated with predicted older non-small cell lung carcinoma CCLs, consistent with increased reliance on membrane remodelling and proteostasis pathways in ageing-associated cellular states^1^.

Several age-associated genetic dependencies were detected in specific tumour contexts (**Fig. 4B**). At the level of individual genes, in Acute Myeloid Leukemia, DDI2 showed a positive correlation with predicted age, indicating reduced dependency in biologically older CCLs. DDI2 encodes a proteasome-associated aspartyl protease involved in activation of the transcription factor NRF1 during proteasome stress responses^49^. Its stronger dependency in younger AML models may therefore reflect a higher reliance on proteasome homeostasis pathways in proliferative leukemic states.

In Bladder Carcinoma, FAM83D and BNIP3 displayed positive correlations with predicted age, indicating reduced dependency in older cell lines. FAM83D is a mitotic spindle-associated protein implicated in cell cycle progression and tumor proliferation. Its high expression has been associated with poor prognosis in various cancers among which bladder urothelial carcinoma^50^. BNIP3 is a hypoxia-inducible regulator of mitochondrial autophagy and metabolic stress response known for providing bladder cancer progression and modulating chemosensitivity as a downstream player of the Ras signaling pathway^51,52^. The reduced dependency observed in older predicted states may reflect shifts away from proliferative mitotic programmes or hypoxia-driven mitochondrial turnover.

By contrast, MEMO1 showed a negative correlation with predicted age, indicating increased dependency in older cell lines. MEMO1 is a mediator of receptor tyrosine kinase signalling and cell motility downstream of ERBB2 and IGF1R pathways, shown to promote bladder cancer cell proliferation and motility in vitro^53^. Moreover MEMO1 is observed to regulate iron homeostasis in cancer through its iron binding capacity^54^. Increased dependency on MEMO1 in older predicted states may therefore reflect compensatory reliance on growth-factor signalling pathways as well as a limited cell motility and iron utilization.

Finally, in the “Other Solid” cancer group, SLC30A9 exhibited a negative correlation with predicted age, suggesting greater dependency in biologically older cell lines. SLC30A9 encodes a mitochondrial zinc transporter involved in maintaining mitochondrial function and redox homeostasis^55^. Increased dependency on this gene in older predicted states is consistent with the broader enrichment of mitochondrial maintenance programmes observed across tumour types, suggesting that ageing-associated epigenetic states may coincide with increased reliance on mitochondrial homeostasis pathways.

Together, these findings highlight tumour-context-specific genetic dependencies associated with ageing-related epigenetic states, supporting the notion that the epigenetic ageing axis captured by GepiClock stratifies functional vulnerabilities across cancer models

## Discussion

Epigenetic clocks (epiclocks) have so far demonstrated accuracy in predicting chronological age of healthy tissues, but their utility in cancer has remained limited because tumour-associated methylation reprogramming disrupts canonical ageing trajectories. Here, we addressed this gap by developing a generalised epiclock (GepiClock), a tumour-trained epiclock that predicts age robustly in primary cancers and, strikingly, generalises to matched normal tissues. By training directly on malignant methylomes across 32 TCGA cancer types, our model captures ageing-associated methylation structure that remains detectable despite cancer-driven epigenetic disruption, enabling a practical ageing-informed readout in oncologic settings.

Beyond predictive performance, GepiClock offers a framework to integrate an ageing-related epigenetic axis into functional cancer genomic data. The model uses a larger CpG set than many established clocks, reflecting both modern array coverage and the heterogeneity introduced by multi-cancer training. Importantly, we show that GepiClock remains accurate under substantial probe missingness and when constrained to CpG set sizes comparable to prior clocks, arguing that its performance does not simply reflect increased model complexity and supporting portability across methylation platforms.

A key conceptual contribution of this work is the use of predicted age in cancer cell lines (CCLs) not as a surrogate for an ill-defined “chronological” CCL age, but as a projection onto tumour-derived ageing-associated methylation states. Despite known epigenomic drift and culture-induced reprogramming in vitro, this projection retained sufficient biological signal to stratify therapeutic vulnerabilities. At the drug-class level, predicted younger CCLs preferentially responded to mTOR, MEK1/2, and HSP90 inhibition, whereas predicted older CCLs showed greater sensitivity to AKT and PI3Kα inhibitors. These associations map onto nutrient-sensing, growth, proteostasis, and survival signalling modules widely implicated in ageing biology, suggesting that GepiClock captures an epigenetic state linked to differential reliance on proliferative versus stress-adaptive programmes. Cancer-type–specific analyses further revealed both recurrent and lineage-restricted enrichments (e.g., EGFR/PI3Kα/BCL2 in older-predicted states versus KIT/MEK in younger-predicted states), indicating that ageing-associated epigenetic states interact with tumour-specific signalling architecture rather than producing uniform pan-cancer effects.

Extending the same framework to genetic perturbation data, GepiClock stratified dependency landscapes in ways consistent with central ageing-related biology. Pathway-level enrichments highlighted mitochondrial gene expression/translation and respiratory processes across multiple tumour types, and we identified tumour-context–specific dependencies associated with predicted age, including DDI2 in AML and MEMO1/SLC30A9 in other settings. Together, these results support the idea that ageing-associated epigenomic organisation can help explain variability in both drug sensitivity and genetic essentiality, providing an orthogonal dimension to standard genomic and transcriptomic stratification.

This study has limitations. Associations derived from cell lines may not fully translate to patient tumours given culture-driven drift, and dose-response measures can be constrained by experimental concentration ranges for some compounds. Moreover, our analyses are observational and do not establish causality between ageing-associated methylation states and dependencies. Future work should validate these findings in patient-derived models and primary tumour cohorts with matched functional measurements, and dissect mechanisms connecting ageing-associated methylation states to pathway wiring and therapeutic response. Despite these caveats, GepiClock provides a broadly applicable tool for studying ageing in cancer contexts and a principled route to discover age-linked therapeutic vulnerabilities.

By enabling ageing-aware stratification of cancer functional data, GepiClock opens the door to precision oncology strategies that explicitly account for the ageing state of tumour cells.

## Methods

### Data sources

We obtained DNA methylation data generated using the Illumina Infinium HumanMethylation450 BeadChip array from TCGA (v38.0, August 31, 2023) through the TCGAbiolinks R package (version 2.30.0), together with sample-level clinical annotations, including age at diagnosis, sample type, sex, and ancestry. We retained only primary tumour, solid tissue normal, and primary blood cancer samples with available age at diagnosis. The final TCGA dataset comprised 9,105 samples across 32 projects, each corresponding to a distinct cancer type (**Supplementary Table 1**).

Although these samples were profiled using the HumanMethylation450 array, we restricted the analysis to CpG sites shared with the Illumina Infinium HumanMethylationEPIC and EPIC v2.0 arrays to maximise cross-platform compatibility. In total, we retained 369,845 CpG probes (**Supplementary Fig. 3**).

The further independent dataset consisted of DNA methylation data of 289 women in the NCI-Maryland Breast Cancer Cohort. It was processed and utilized for model performance validation^33^. It includes 185 tumor samples, 113 additional paired adjacent normal samples, and 104 normal tissues from reduction mammoplasty (GEO accession number GSE225845).

We obtained cancer cell line (CCLs) methylation data from GEO (accession GSE68379)^23^. This dataset includes 1,028 human cancer cell lines profiled using the HumanMethylation450 array, together with metadata such as cancer type, tissue of origin, molecular subtype, and age at sampling (available for 690 cell lines).

To harmonise CCL annotations and integrate multiple data sources, we used the Cell Model Passports (CMP) platform (model list 20240110). We extracted cancer type, tissue of origin, and microsatellite instability (MSI) status, and used identifiers (BROAD_ID, Sanger_ID, and CAccession) to integrate data across DepMap and GDSC.

We obtained drug response data from the Genomics of Drug Sensitivity in Cancer (GDSC) datasets (https://www.cancerrxgene.org/, GDSC1 and GDSC2, fitted dose–response v27Oct23)^40^, which quantify the sensitivity of cancer cell lines to a broad range of chemotherapeutic and targeted agents. We integrated log(IC50) values into our analysis pipeline alongside genomic features, including mutations, copy number alterations, and gene expression profiles, to assess the relationship between methylation patterns and drug response.

Finally, we obtained CRISPR-based gene effect scores from the Cancer Dependency Map (https://depmap.org/, Public release 24Q2)^22^, which provide a continuous measure of gene essentiality across cancer cell lines. We used these scores together with reference sets of common essential and non-essential genes from the Achilles dataset for downstream analyses.

### Methylation data pre-processing and normalization

We processed raw DNA methylation data in IDAT format using the minfi R package (version 1.48.0). We performed background correction using the single-sample normal-exponential out-of-band (ssNoob) method.

To assess data quality, we computed detection p-values for each CpG probe. We set probes with p-values > 0.05 to missing (NA) and excluded samples with more than 10% missing CpG values. We further removed cross-reactive probes, CpH probes and probes located on sex chromosomes (chrX and chrY).

We imputed the remaining missing values using the k-nearest neighbours (KNN) algorithm implemented in the ChAMP R package (version 2.8.6). We then applied Beta Mixture Quantile (BMIQ) normalisation, also using ChAMP, to correct for probe-type bias inherent to the array design.

We report the number of samples and CpGs removed, as well as the number of imputed values at each processing step, in **Supplementary Table 1**.

### Elastic net model

We trained an elastic net regression model to predict age from DNA methylation data using the glmnet (version 4.1-8) and glmnetUtils (version 1.1.9) R packages. We performed cross-validation to identify optimal values for the regularisation parameters α and λ.

We split TCGA data into training, test, and evaluation sets (**Fig. 1A**). For model training and testing, we retained only primary tumour samples from unique patients to avoid donor-related bias. We partitioned the data into 80% training and 20% test sets. To preserve representation across cancer types, we performed the split within each TCGA project. We further ensured that age distributions remained comparable between training and test sets within each cancer type using the caret R package (version 6.0.94). We reserved samples from patients with multiple specimens for independent evaluation of model performance across matched tumour and normal samples.

We trained the model on 6,142 tumour samples spanning 32 cancer types. To ensure compatibility with newer methylation platforms, we restricted the analysis to CpG probes shared across the Illumina HumanMethylation450, EPIC, and EPIC v2.0 arrays, thereby avoiding missing values in downstream applications. We used the cva.glmnet function to perform joint cross-validation of α and λ using 10-fold cross-validation.

We selected the final model based on the combination of α and λ that minimised the mean squared error (MSE). For each α, we used the λ.1se criterion, defined as the largest λ within one standard error of the minimum cross-validated error (λ.min), to favour model generalisability.

We evaluated model performance using MSE, root mean squared error (RMSE), and Pearson correlation (R). We assessed predictive accuracy on independent datasets not used for training, including both tumour and matched normal samples across TCGA projects, as well as on the external independent dataset^33^.

### Randomly subsampled GepiClock

To evaluate the robustness of GepiClock to missing values, we performed repeated random subsampling of predictive CpGs (20 iterations per subsampling level). At each level, we retained a decreasing portion of predictive CpGs and evaluated model performance on tumour samples from the test set, as well as on both tumour and normal samples from the independent evaluation set.

This analysis allowed us to assess model performance under progressively reduced feature availability and to simulate scenarios with incomplete CpG measurements. It also enabled direct comparison with existing epiclocks by restricting GepiClock to the same number of predictive CpGs used in those models, thereby controlling for feature set size.

### Sample stratification for tissue- and cancer-type analyses

To ensure robust statistical modelling and avoid biases arising from underrepresented groups, we implemented a threshold-based classification strategy to determine whether to use tissue type or cancer type as the primary factor in the pan-cancer ANOVA model and as grouping variables in cancer type-specific analyses. This strategy was necessary because cell line annotations in the Cell Model Passports (CMP) are hierarchically structured, with cancer types nested within broader tissue categories. As a result, the appropriate level of stratification depends on the number of cell lines available within each group.

We therefore based the classification on group sample size. Specifically, we retained tissues with at least 10 associated cell lines as independent categories, while we grouped tissues with fewer than 10 cell lines into an “other” category. For tissues meeting this threshold, we further evaluated the distribution of cell lines at the cancer-type level. When at least one cancer type within a given tissue included 10 or more cell lines, we used cancer type as the primary stratification variable; otherwise, we retained tissue type as the grouping factor (**Supplementary Fig. 12A**).

Applying this procedure resulted in the classification of 864 cell lines for the drug response analysis and 561 for the gene dependency analysis. The majority of cell lines were stratified by cancer type; however, less represented cancer types were grouped at the tissue level, particularly for Bone and Haematopoietic and Lymphoid tissues. Cell lines that did not meet the minimum threshold at either level were assigned to the “other” category.

For cancer type-specific analyses, we further restricted the dataset to cancer types with at least 10 cell lines to ensure sufficient statistical power.

### ANOVA model

To ensure robust statistical modelling and avoid biases arising from underrepresented groups, accounting for biological and experimental variability across cancer cell lines, we applied an analysis of variance (ANOVA) framework to isolate the contribution of predicted biological age. This approach allowed us to estimate and remove the effects of tissue/cancer type and microsatellite instability (MSI) status on drug response or genetic dependency, thereby enabling analysis of age-associated signals independent of these confounding factors. We followed a strategy similar to that described in our previous work^23^.

We implemented the ANOVA model using the base R function aov(). The response variable was either the natural logarithm of IC50 values (ln(IC50)), representing drug sensitivity, or gene effect scores, representing gene essentiality. We included two categorical factors: (i) tissue of origin or cancer type (as determined by the stratification procedure) and (ii) MSI status. The model can be written as:

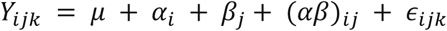

Where:

– *Y_ijk_* denotes the observed ln(IC50) or gene effect value;
– *μ* is the overall mean
– *α_i_* represents the effect of tissue or cancer type
– *β_j_* the effect of MSI status
– (*αβ*)*_ij_* their interaction and
– *∈_ijk_* the residual error.

After fitting the model, we extracted residuals for each cell line, representing variation not explained by the included factors. Residuals satisfied normality assumptions (**Supplementary Fig. 14**). We then correlated these residuals with predicted biological age to quantify age-associated effects on drug response and genetic dependency. For cancer type-specific analyses, we simplified the model by including only MSI status when variability was present. In the absence of MSI heterogeneity, we directly computed correlations without ANOVA adjustment.

We generated all plots using *ggplot2* (v3.4.4). For enrichment visualisations (**Fig.s 3C** and **4A**), we applied the *MutExMatSorting* package (v0.1.0)^56^ to reorder matrices and improve interpretability by grouping partially overlapping patterns.

### Genetic Dependency Data pre-processing

We obtained genetic dependency data from the DepMap Public 24Q2 release, using CRISPR gene effect scores corrected with the Chronos algorithm. Chronos models cell proliferation dynamics following CRISPR-mediated gene knockout and corrects for known biases, including copy number and proximity effects, as well as screening quality and DNA damage-related noise^57,58^.

We retrieved lists of common essential and non-essential control genes (Achilles datasets) from the same release and excluded them from the analysis. We further scaled gene effect data relative to essential and non-essential gene sets and applied the CoRe.FiPer function from the CoRe package^59^ to identify core fitness genes using the area under the curve (AUC) method. We classified genes with scores above the 90th percentile as core fitness genes and removed them from subsequent analyses.

We extended this filtering by excluding additional essential and non-essential gene sets from the BAGEL dataset, including curated essential genes involved in fundamental cellular processes such as DNA replication, histone formation, ribosomal function, proteasome activity, and RNA polymerase complexes^60^. This step ensured that downstream analyses focused on genes with variable dependency profiles across cell lines.

We further filtered genes by requiring a median scaled depletion effect of at least −0.5 in at least two cell lines. After applying all filters, the gene effect matrix comprised 6,052 genes out of the original 18,443.

We applied the same filtering strategy in cancer type-specific analyses. However, because CoRe.FiPer identifies core fitness genes in a context-dependent manner, the number of retained genes varied across cancer types, reflecting lineage-specific dependency patterns.

### Gene and Drug Set Enrichment Analysis

For the gene set enrichment analysis (GSEA) we used as query signatures terms from Gene Ontology^61^ for Biological Process (GOBP), Molecular Function (GOMF), Cellular Component (GOCC) as well as Kyoto Encyclopedia of Genes and Genomes (KEGG)^62^. For the drug set enrichment analysis we used as query signatures, a set of drugs targeting the same protein according to the drug annotation available on the GDSC data portal. A pan-cancer and cancer type specific GSEA and DSEA was then performed (separately for each group of cell lines belonging to the same cancer type or at the pancancer setting) using as background ranked list all genes sorted based on the Pearson correlation of their essentiality score and predicted age across cell lines (for GSEA) or all drugs sorted based on the Pearson correlation of their ln(IC50) and predicted ages across cell lines (for DSEA).

## Data availability

All the data used in this study is publicly available and accessible as described in the relevant Methods’ sections.

## Code availability

All code required to reproduce the analyses, results, and figures presented in this manuscript is available at: https://github.com/ireneefr/GepiClock

## Acknowledgements

This work was partially supported by an AIRC (Associazione Italiana Ricerca sul Cancro) Investigator Grant to FI (code: 28772).

Irene Fernández-Rebollo is a PhD student within the European School of Molecular Medicine (SEMM).

## Author Contributions

IFR, AD, ME and FI conceived the project and scope; IFR, AD, and AO collected data and metadata; IFR and AD, processed and analysed data; IFR designed and implemented the GepiClock pipeline; AD implemented and conducted the drug-response/genetic-dependency association analyses; IFR, AD, AO and FI drafted the manuscript and designed the figures; IFR, AD, AO, LT, FI and ME edited and revised manuscript and figures; AO and ME contributed to study supervision; FI supervised the study.

## Declaration of generative AI and AI-assisted technologies in the writing process

During the preparation of this work, the authors used ChatGPT (OpenAI) to assist with language editing and proofreading, with the aim of improving clarity, grammar, and readability. No content generation, data analysis, or scientific interpretation was performed by the tool. All authors reviewed, revised, and approved the final version of the manuscript and take full responsibility for its content.

