## Supplementary Material for "A Generalised Epigenetic Clock Reveals Therapeutic Vulnerabilities Linked to Ageing in Cancer Cells"

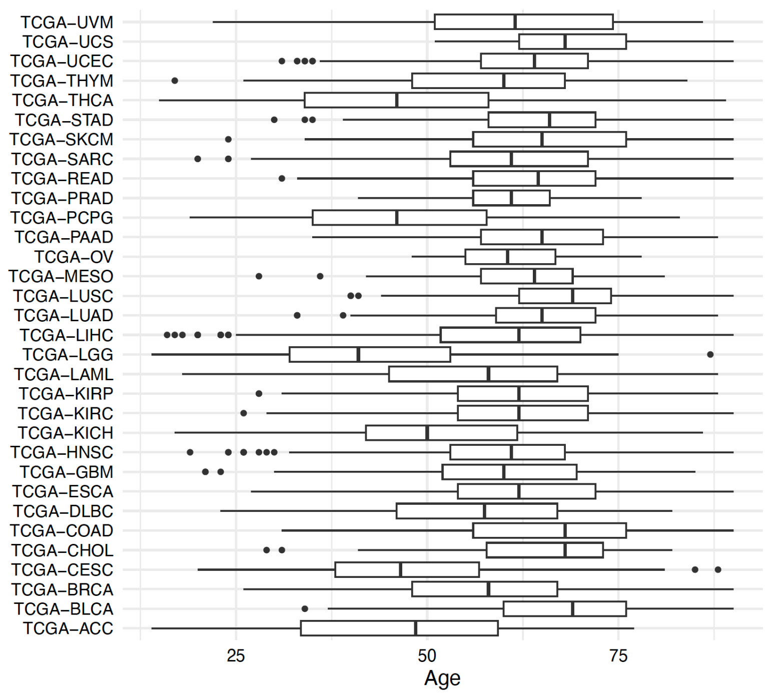


**Supplementary Figure 1 - Chronological age distribution across TCGA projects considered in the study.**

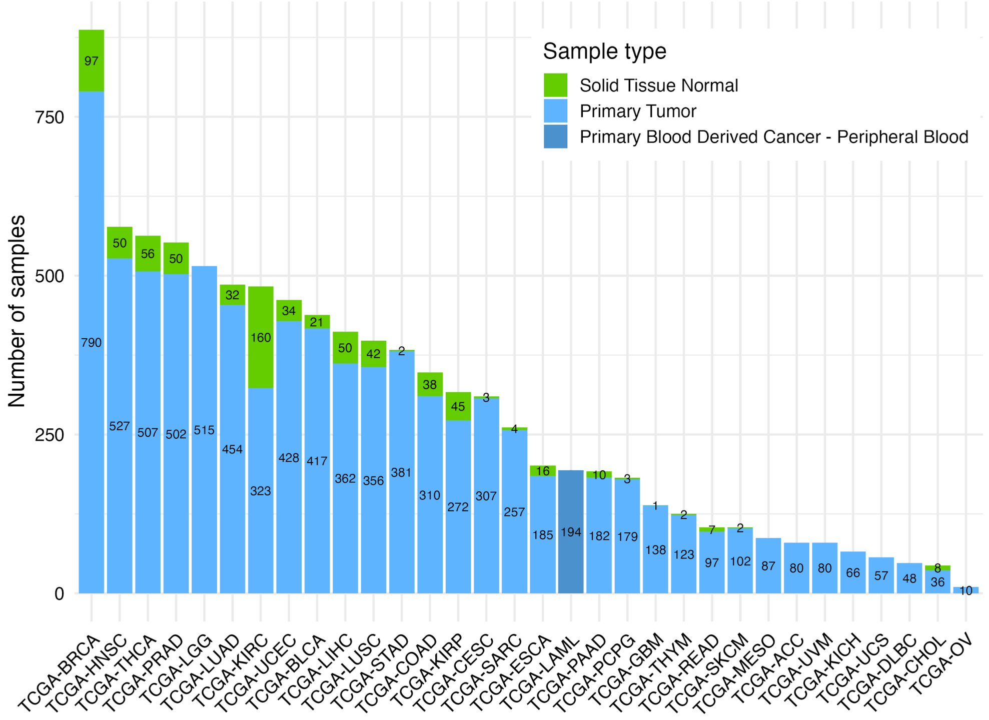


**Supplementary Figure 2 - Number of Samples across TCGA projects included in our analysis.**

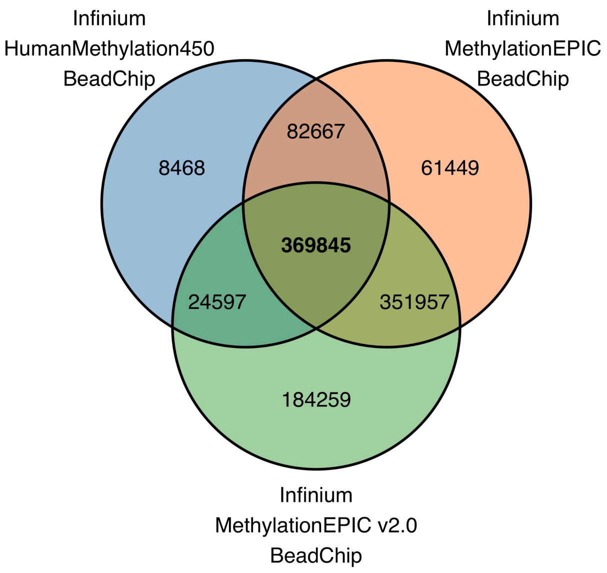


**Supplementary Figure 3 - Number of CpGs shared across different Illumina Infinium Methylation Beadchips.**

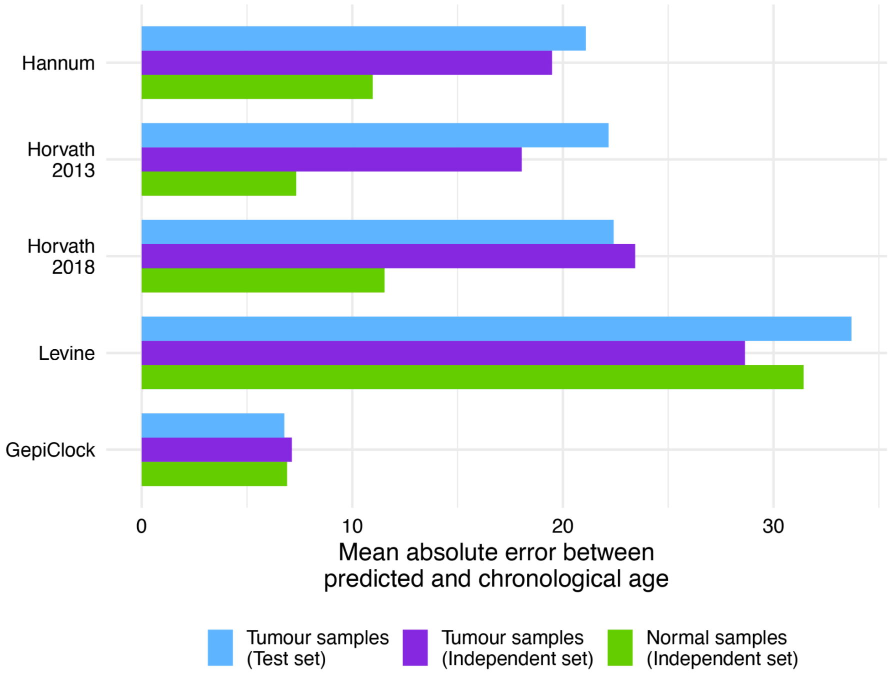


**Supplementary Figure 4 - Mean absolute error between predicted and chronological age of widely used epigenetic clocks and of the GepiClock across different data type subgroups (Tumour samples in the test set, tumour samples in the held-out independent set, and normal samples in the held-out independent set).**

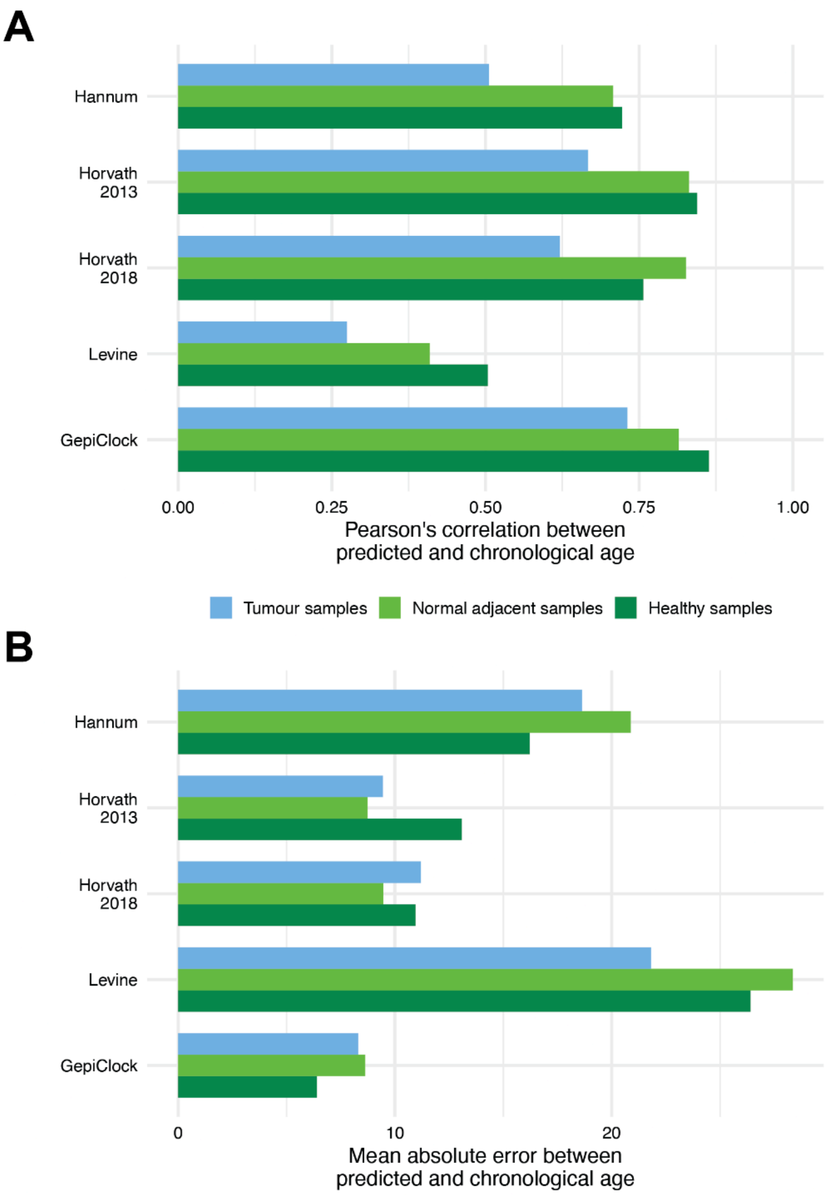


**Supplementary Figure 5 - Predictive performances of widely used epigenetic clocks and of the GepiClock across different data type subgroups from the GSE225845 dataset (Tumour samples, normal adjacent samples, and healthy samples).**

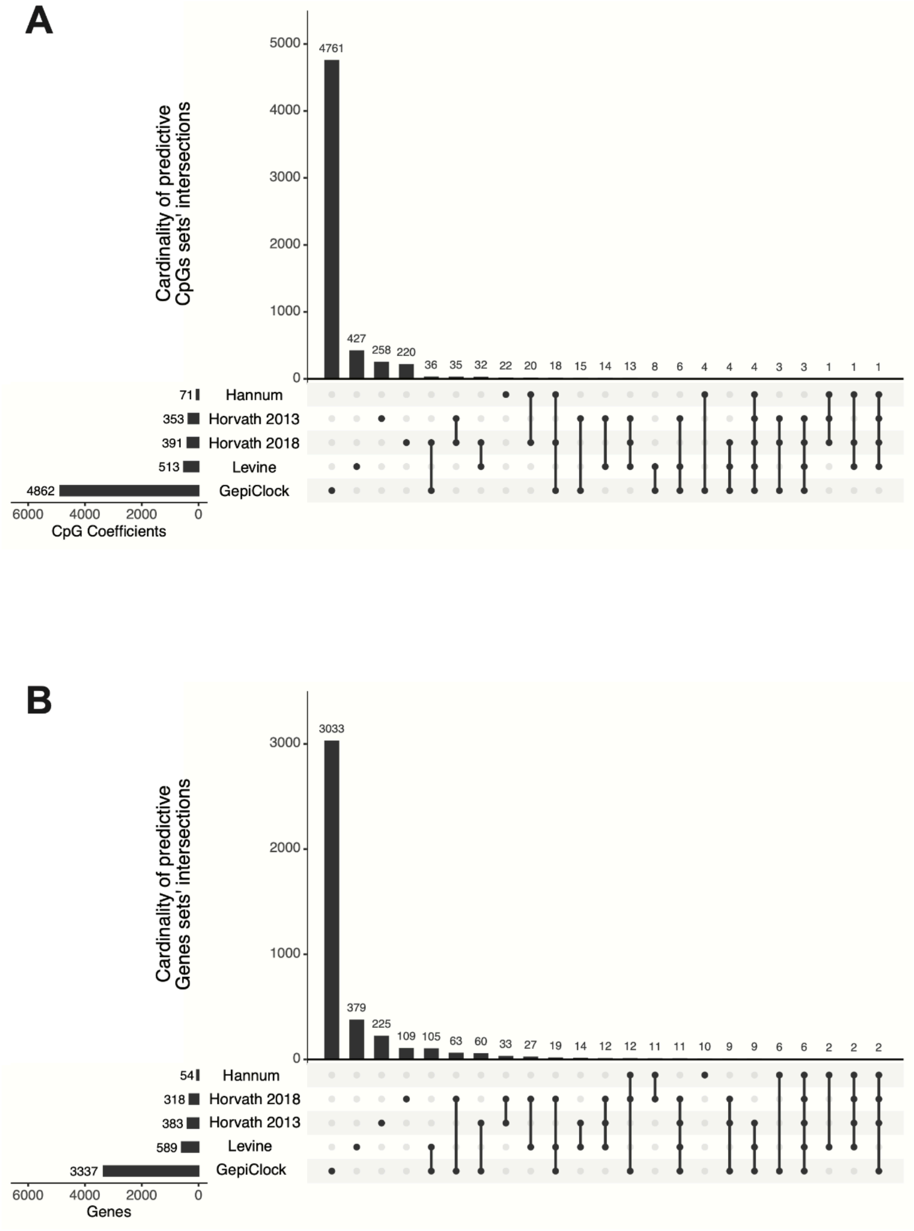


**Supplementary Figure 6 - Number of predictive CpGs (A) and genes (B), and their overlaps, across different epigenetic clocks.**

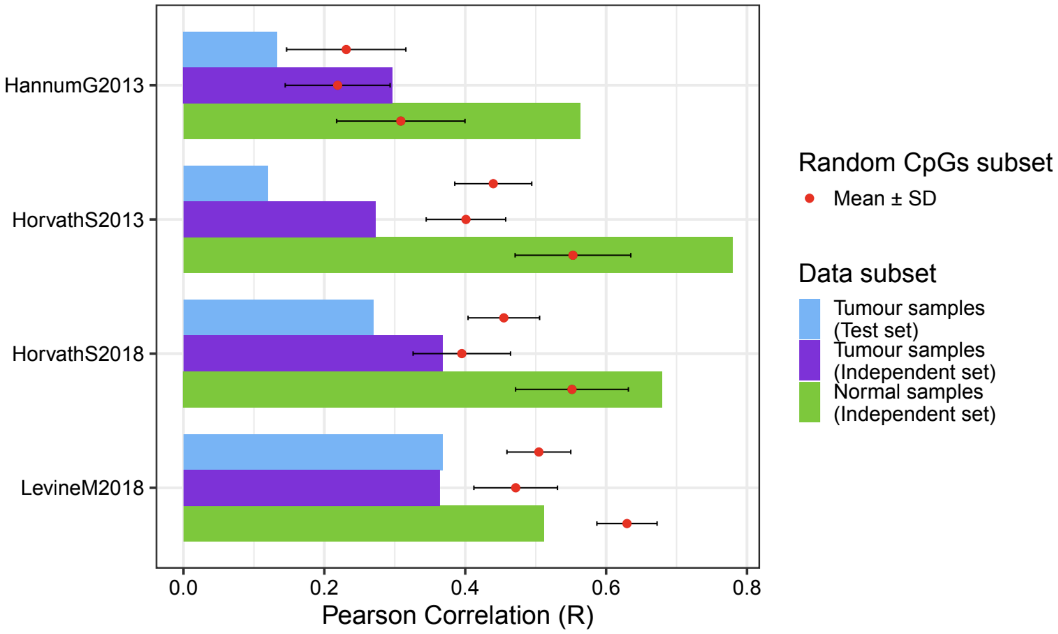


**Supplementary Figure 7 - Epigenetic clocks performance versus the subsampled GepiClock.**
Model-size-matching comparison of the predictive performances (Pearson's correlation of predicted vs chronological age) between the GepiClock and published epiclocks. Bars indicate the performances of the published clocks across different sample subsets (as per the color legend), whereas whisker plots indicate the performances of the GepiClock when keeping in the model only *x* randomly subsampled predictive CpGs, where *x* is equal to the number of the CpGs in the published epiclock under consideration. Average performances are reported across 20 sampling trials (the red dots) with whiskers indicating mean ± standard deviation.

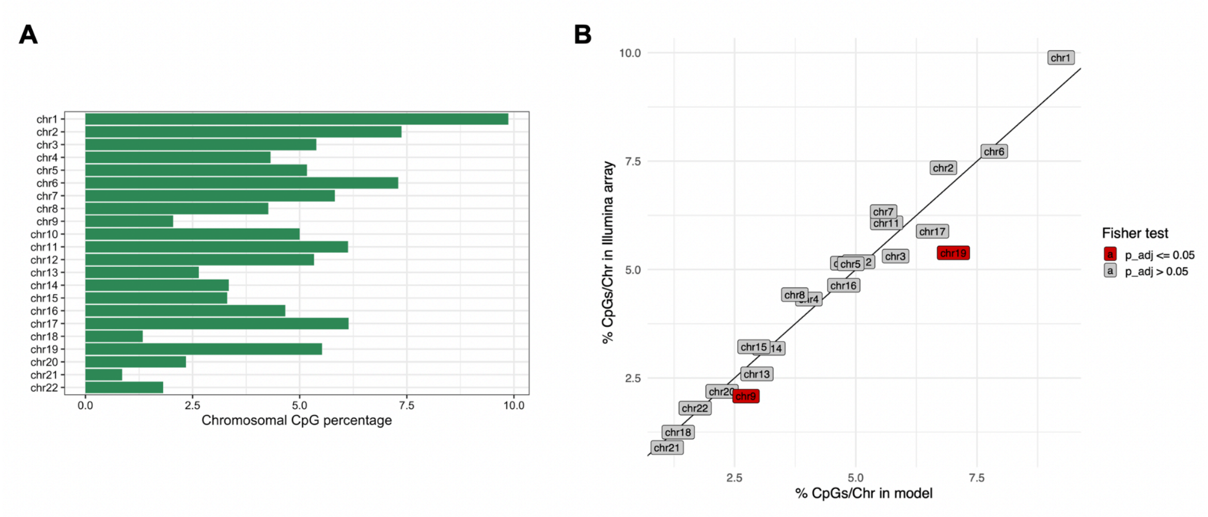


**Supplementary Figure 8 -** A. Percentages of Illumina arrays' CpGs across chromosomes; B. Distribution of CpGs (percentage over total) in the GepiClock (x-axis) versus total number in Illumina Array (y-axis) across autosomal chromosomes (the individual points). Red indicates significant enrichments (Adjusted Fisher exact test p-value < 0.05).

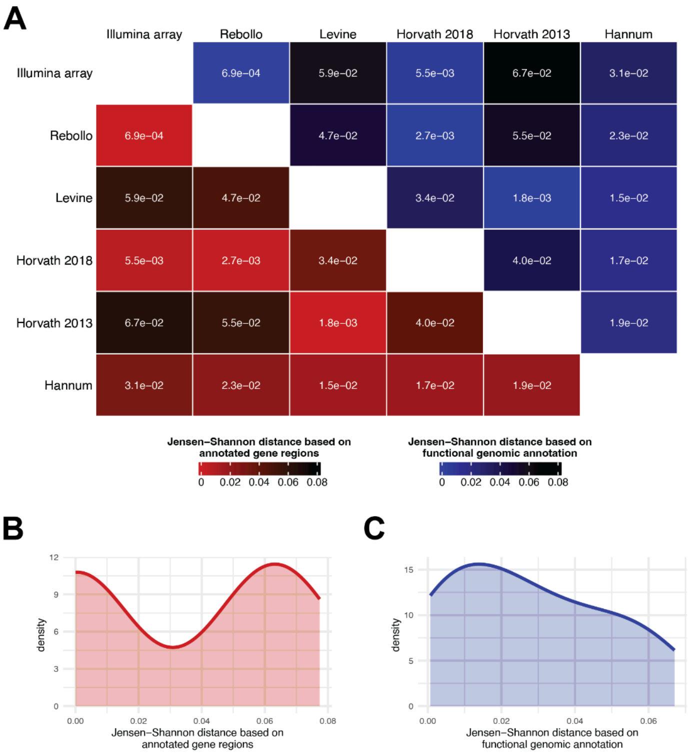


**Supplementary Figure 9 - Epigenetic clocks similarity based on genetic annotations.**

A. Pair-wise Jensen-Shannon distance of annotated/non-annotated gene regions (Gene, No gene, in shades of blue) and functional genomic annotation (TSS1500, TSS200, 5'UTR, 1st Exon, Body, 3'UTR, in shades or red) across predictive CpGs in the different epigenetic clocks and the Illumina 450k array, and related observed values' distributions, shown respectively in B and C.

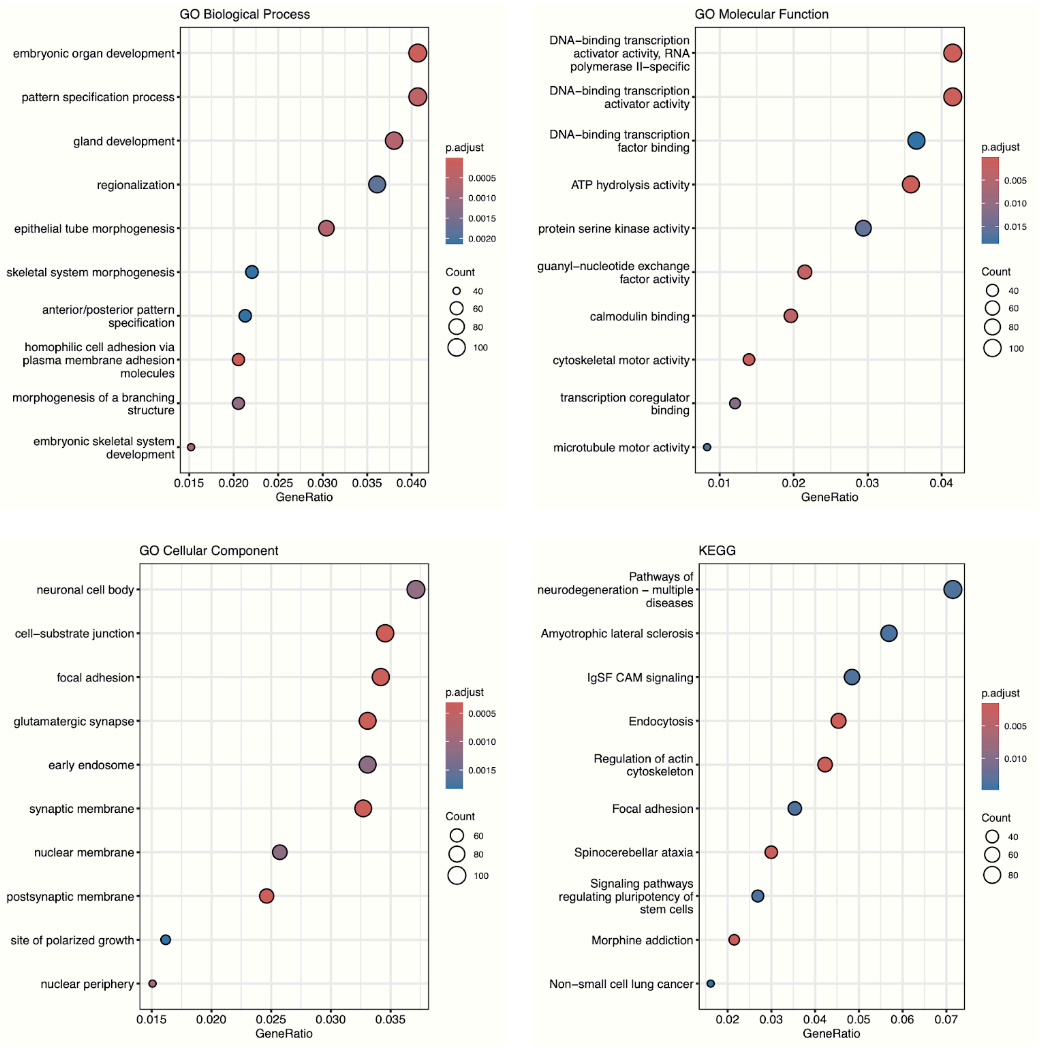


**Supplementary Figure 10 - Functional enrichment analysis of the 3,337 genes in proximity loci of the GepiClock CpGs.** Top 10 enriched categories across gene signature collections are shown. Full list of enrichments is available in Supplementary Table 3.

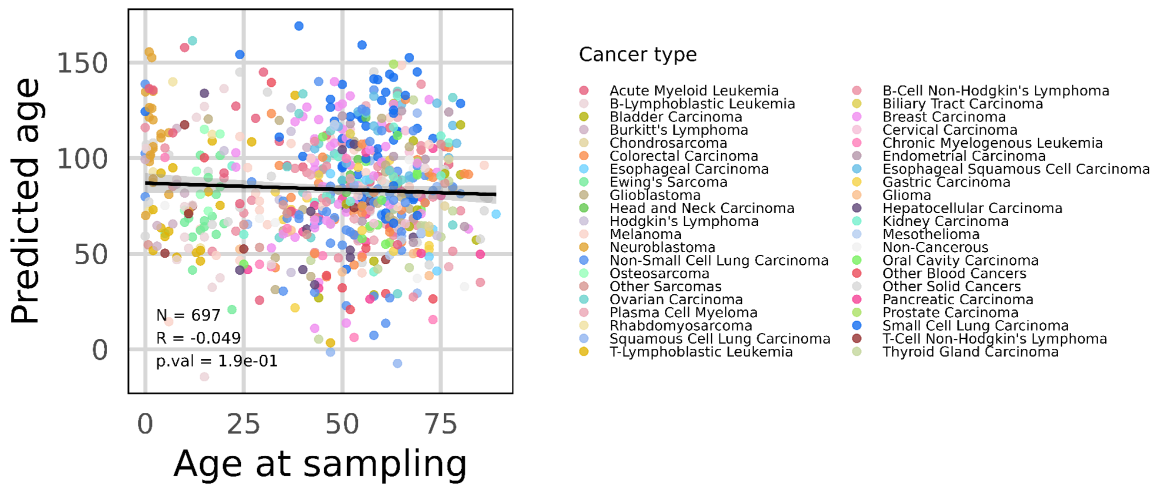


**Supplementary Figure 11 - Lack of correlation between cell lines' age at sampling and their GepiClock-predicted age**. Pearson correlation coefficient (R) computed across the 697 cell lines with available age at sampling annotations and methylation data. Each point represents one cell line and is color-coded by cancer type. The correlation is reported together with its statistical significance (p.val).

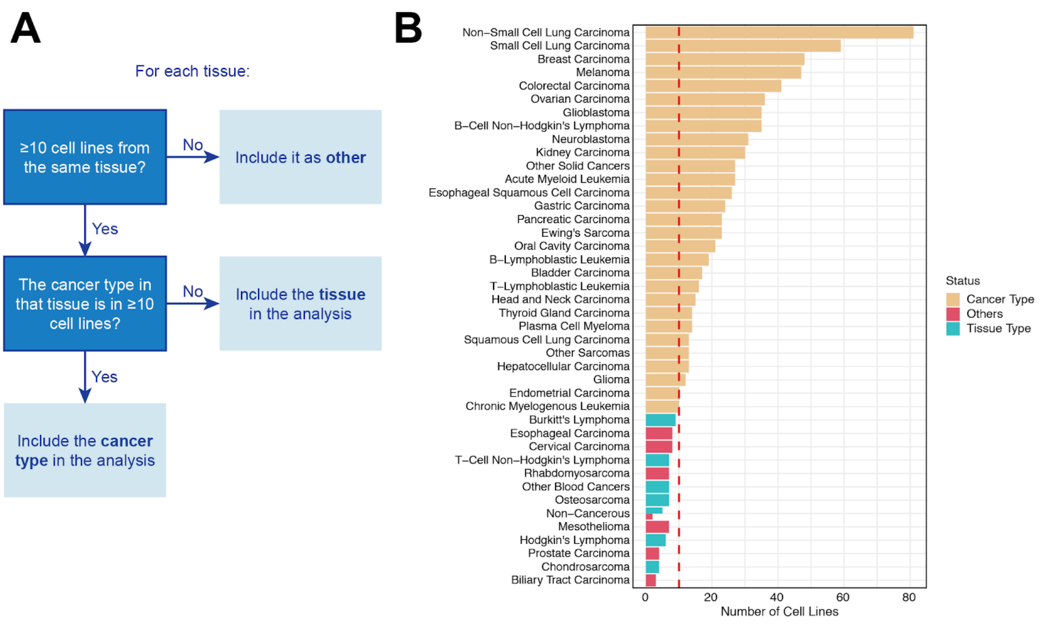


**Supplementary Figure 12 - Cancer Cell Lines Category Selection (drug response analysis).**
**A.** Criteria used to determine cancer- or tissue-type-specific analyses to perform in the association study between predicted cell line age and drug response. **B.** Results of the selection algorithm for the association study between predicted cell line age and drug response. This selection determines whether a cell line should be considered in a tissue- or cancer-type specific analysis, or categorised as "Other".

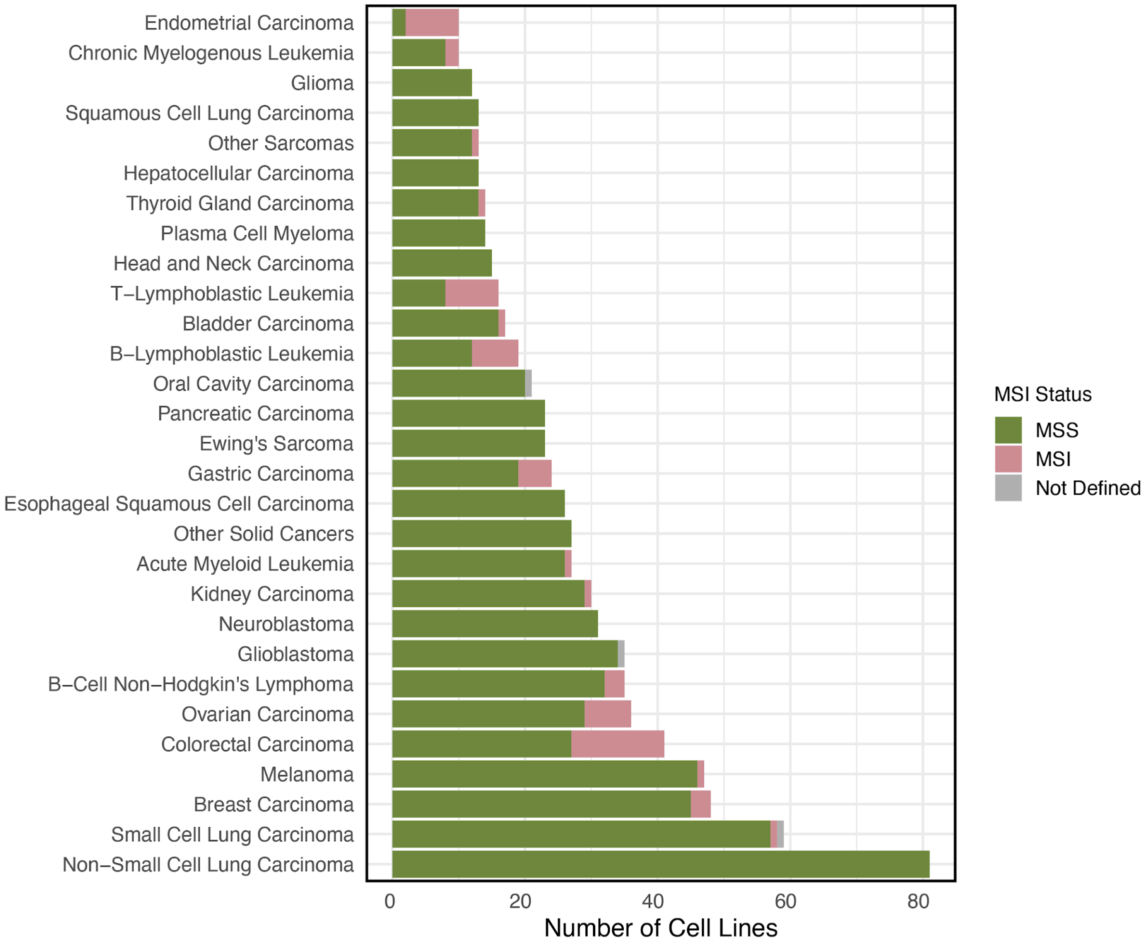


**Supplementary Figure 13 - Cancer Types included in the drug response analysis.**
Distribution of available cell lines across cancer-specific Analysis by Cancer Type and MSI Status. The cancer types with fewer than 10 distinct cell lines were excluded from the association analysis between predicted cell line age and drug response.

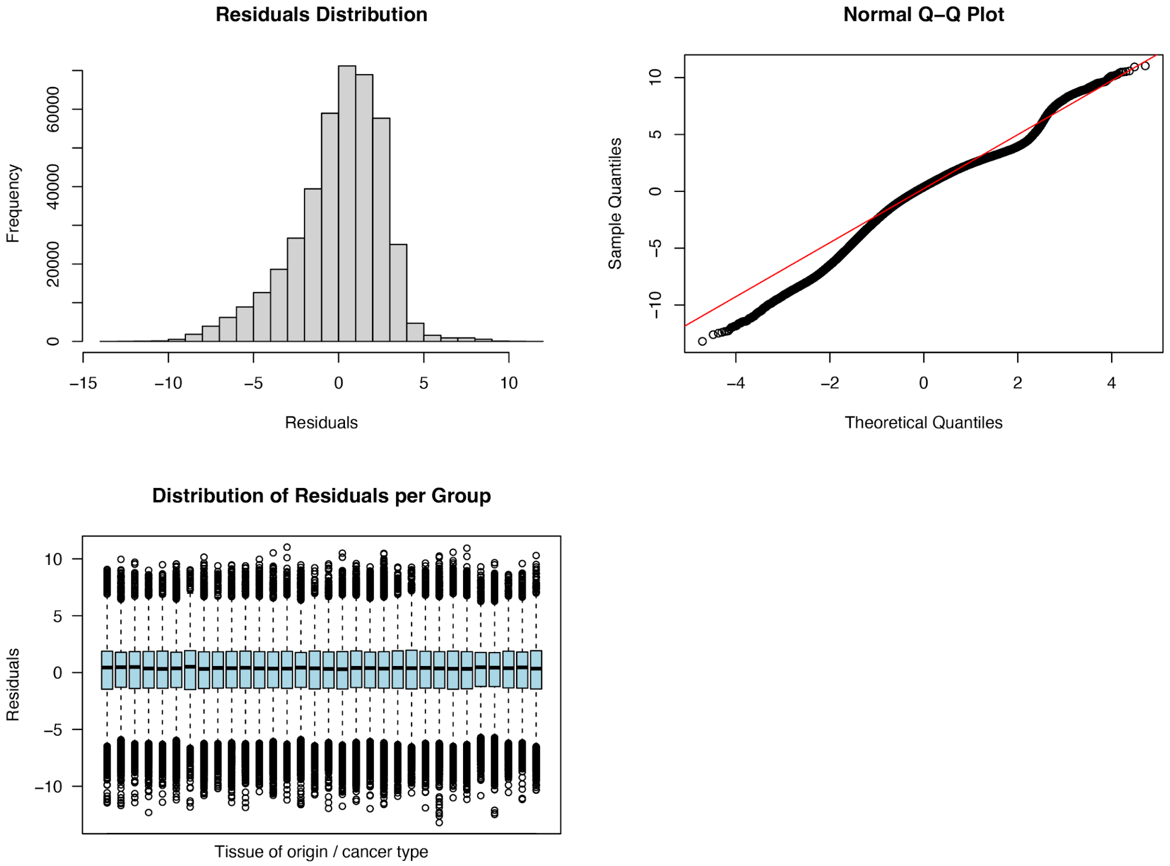


**Supplementary Figure 14 - Normality of drug sensitivity (ln(IC50)) residuals extracted from ANOVA model.**

The distribution of the residuals obtained from the ANOVA model applied to the ln(IC50) values exhibit an approximately symmetrical bell-shaped curve, suggesting a distribution close to normality (S8A). In the Q-Q plot (S8B) most of the points closely follow the red diagonal line, indicating that the residuals are approximately normally distributed. The boxplot (S8C) shows that the residuals are centered around zero for all groups (tissue of origin/cancer type chosen by the selection algorithm) and the spread appears consistent across groups, with no major outliers or systematic patterns observed validating ANOVA model assumptions.

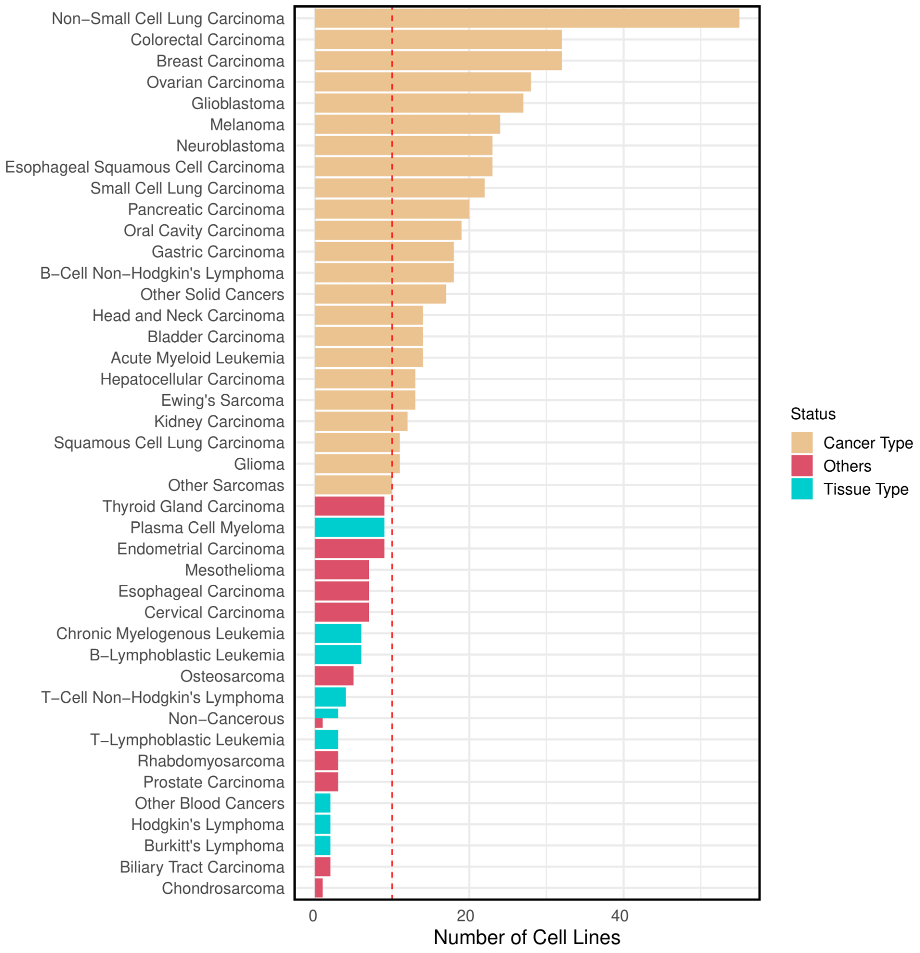


**Supplementary Figure 15 - Cancer Cell Lines Category Selection (gene dependencies analysis).**
Results of the selection algorithm for the association study between predicted cell line age and genetic dependencies. This selection determines whether a cell line should be considered in a tissue- or cancer-type specific analysis, or categorised as "Other", as described in **Supplementary Figure 12.**

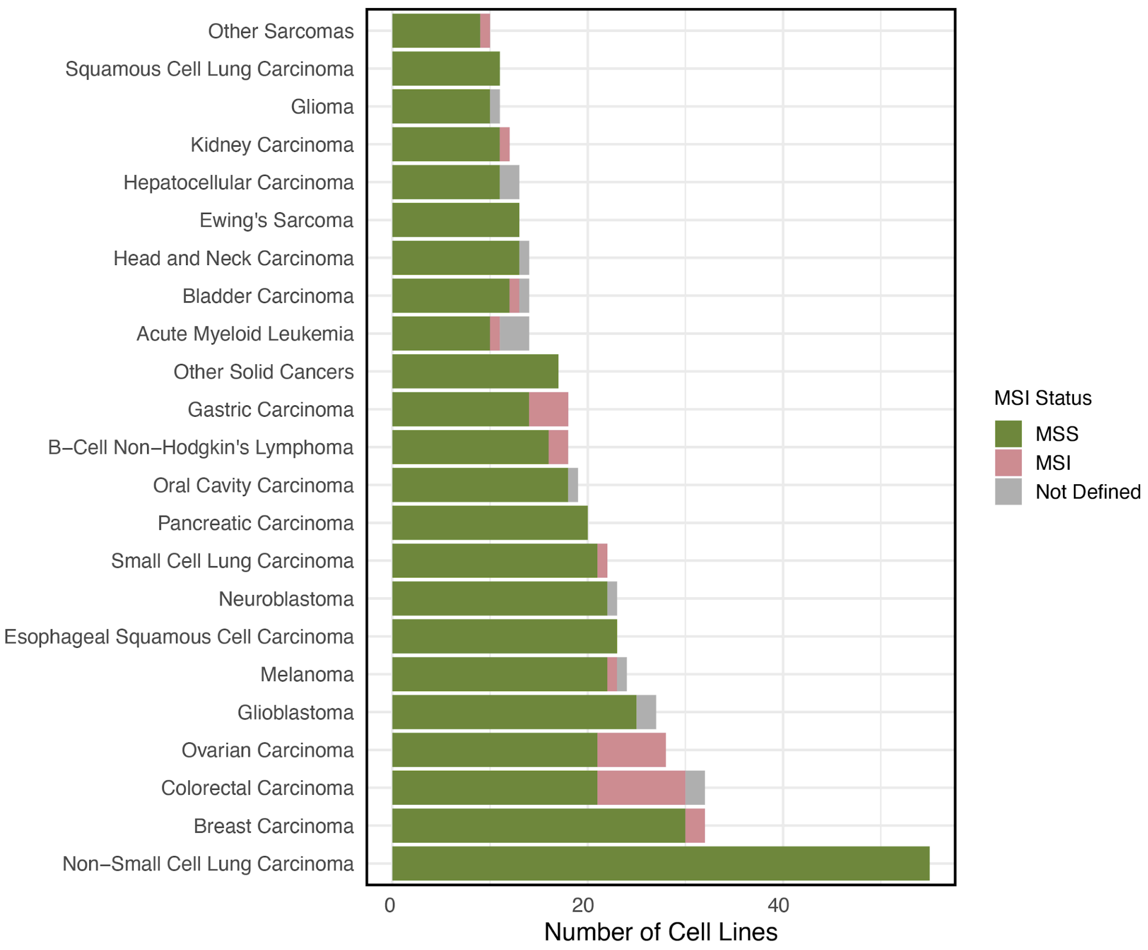


**Supplementary Figure 16 - Cancer Type Selection of Cell Lines (gene dependencies analysis).**
Distribution of Cell Lines in Cancer-Specific Analysis by Cancer Type and MSI Status. The cancer types with fewer than 10 distinct cell lines were excluded from the association analysis between predicted cell line age and genetic dependencies.

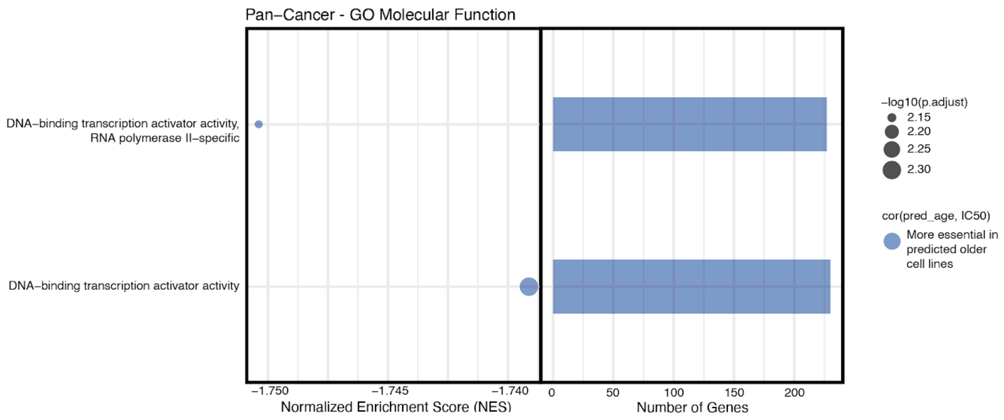


**Supplementary Figure 17 - Gene Set Enrichment Analysis with Gene Ontology Molecular Function terms (GOMF)**

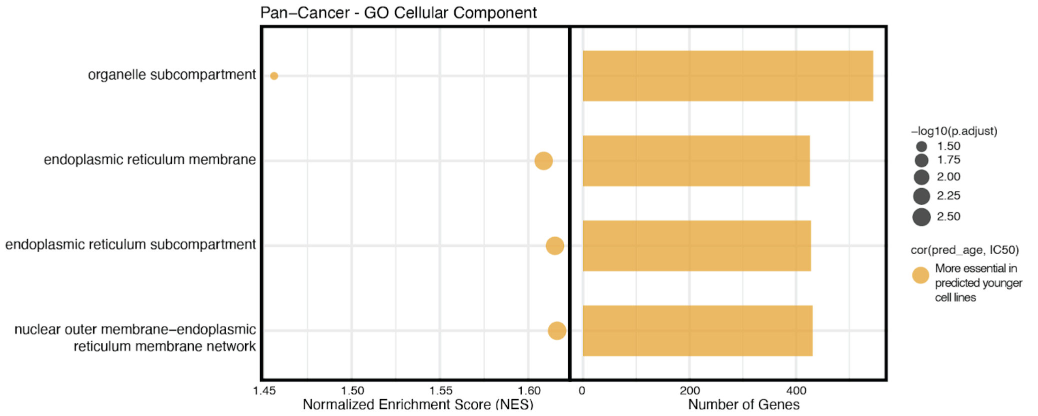


**Supplementary Figure 18 - Gene Set Enrichment Analysis with Gene Ontology Cellular Component terms (GOCC)**

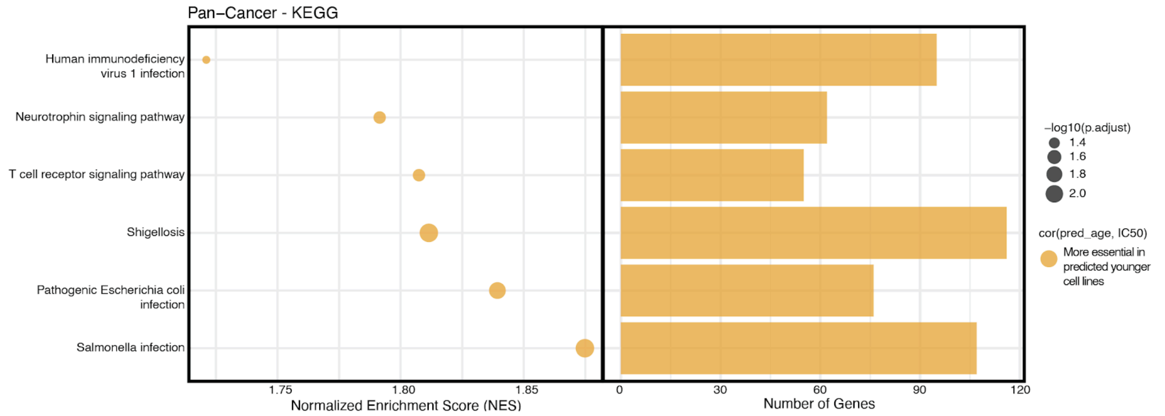


**Supplementary Figure 19 - Gene Set Enrichment Analysis with Kyoto Encyclopedia of Genes and Genomes terms (KEGG)**

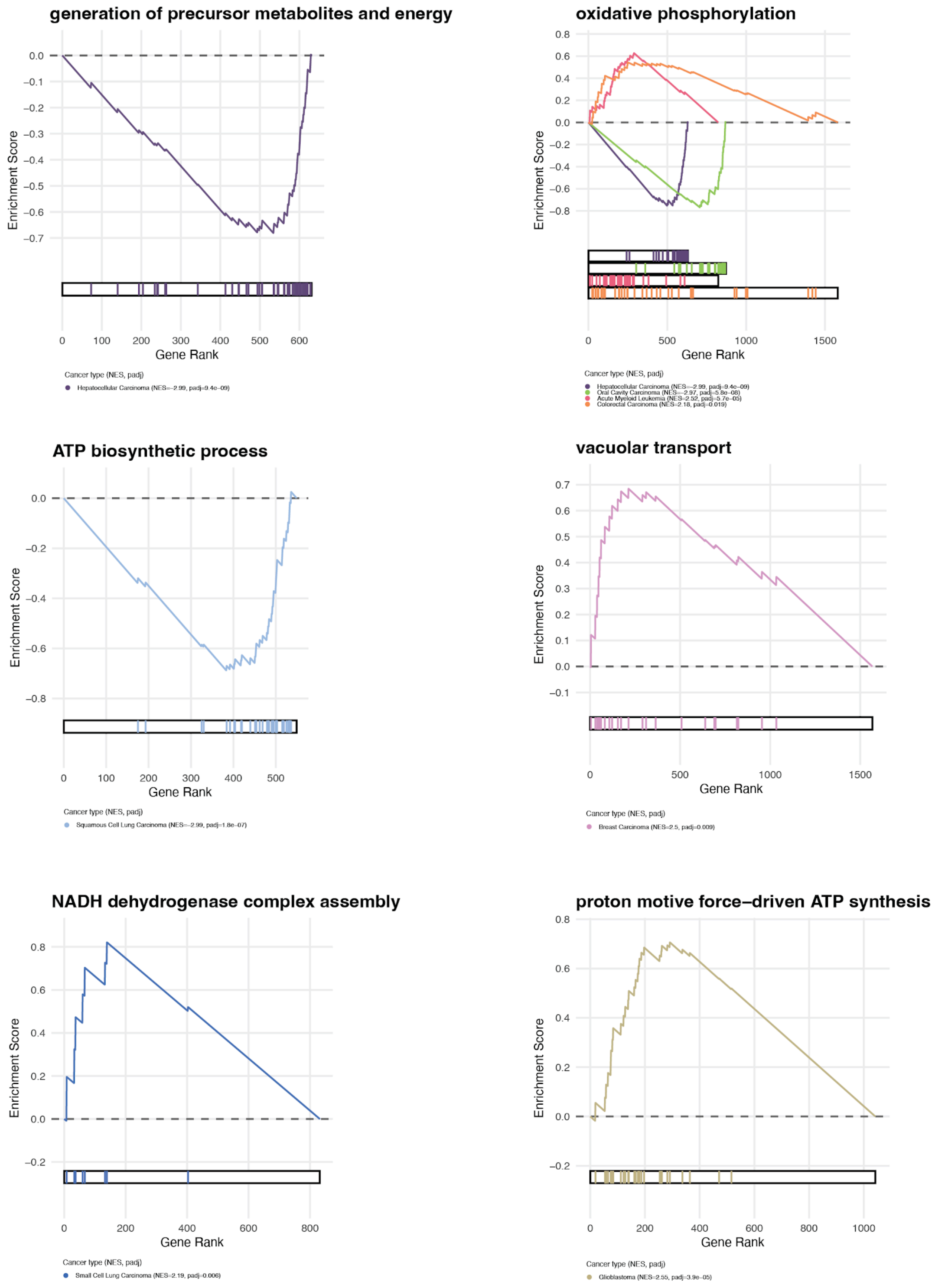


**Supplementary Figure 20 - Cancer type specific Gene Set Enrichment Analysis Significant Results**

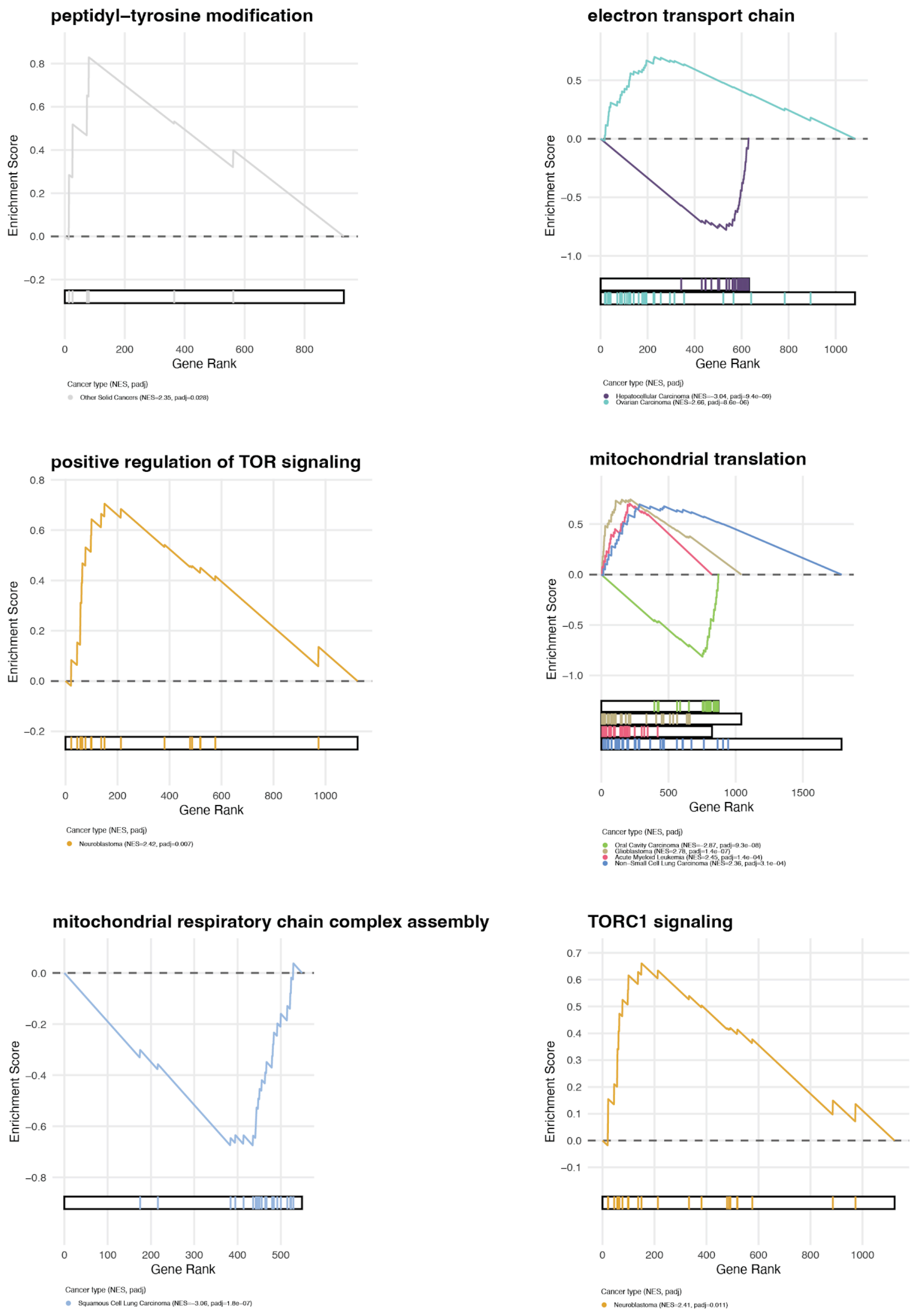


**Supplementary Figure 21 - Cancer type specific Gene Set Enrichment Analysis Significant Results**

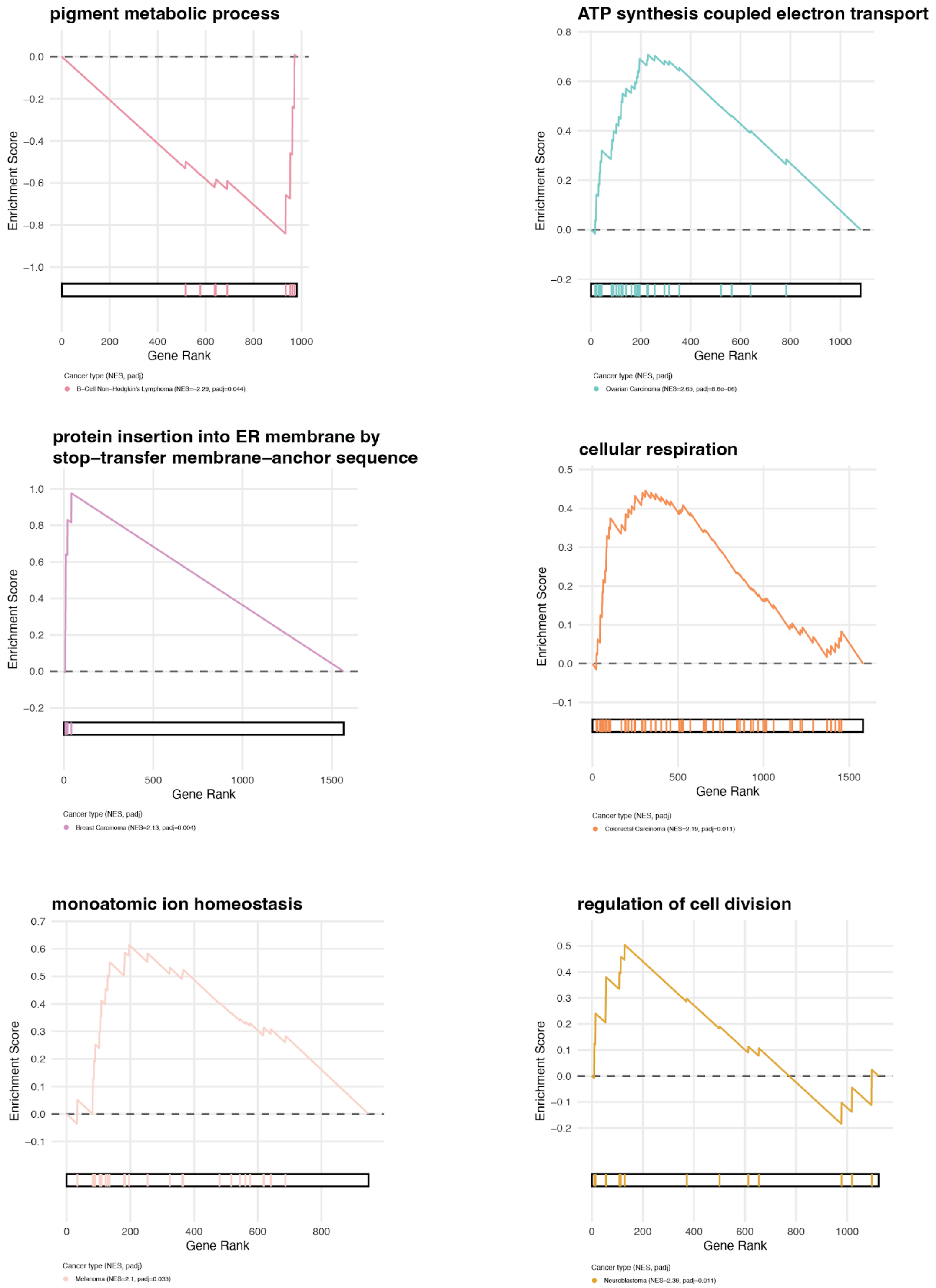


**Supplementary Figure 22 - Cancer type specific Gene Set Enrichment Analysis Significant Results**

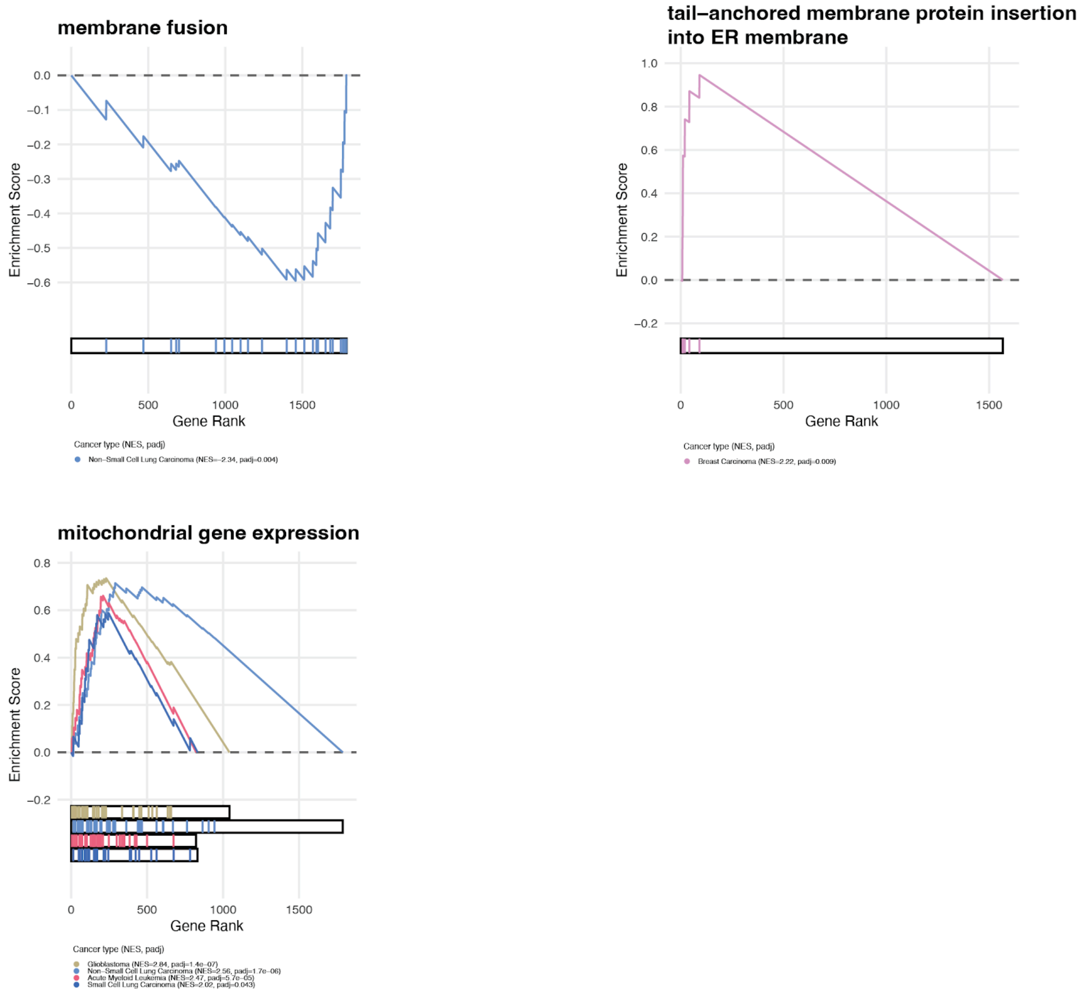


**Supplementary Figure 23 - Cancer type specific Gene Set Enrichment Analysis Significant Results**
